# Mechanism of tandem-repeat DNA synthesis by an antiviral reverse transcriptase

**DOI:** 10.64898/2026.06.29.735271

**Authors:** Josephine L. Ramirez, Qixiang He, Tanner Wiegand, George D. Lampe, Jing Wang, Stephen Tang, Israel S. Fernández, Samuel H. Sternberg

## Abstract

Defense-associated reverse transcriptases (DRTs) employ DNA synthesis to protect bacteria against phage infection^1,2^. We previously showed that DRT10, a tripartite system comprising an RT, a noncoding RNA (ncRNA), and a SLATT effector protein, catalyzes protein-primed, tandem-repeat DNA synthesis in a mechanism strikingly analogous to eukaryotic telomerase^3^. However, the structural basis by which the RT–ncRNA complex directs repeat addition processivity and controls repeat length remains unknown. Here we present cryo-EM reconstructions of two evolutionarily diverse DRT10 RT–ncRNA systems that reveal an unanticipated 2:1 architecture, wherein two RT monomers bind opposite sides of a single, pseudo-symmetric ncRNA. Biochemical experiments demonstrate that each RT monomer reverse transcribes the template encoded on its respective side of the ncRNA, but only one generates the long repetitive product, with the template sequence defined by the distance between two flanking stem-loop anchors. Together with earlier studies of DRT2, DRT3, and DRT9^4–6^, our findings identify a conserved mechanistic logic underlying ncRNA-templated tandem-repeat synthesis across Class 2 UG antiviral systems, despite vastly different architectural solutions.

## INTRODUCTION

Reverse transcriptases (RTs) were first discovered by David Baltimore and Howard Temin within the context of retroviruses, where they catalyze the RNA-dependent synthesis of complementary DNA (cDNA) for integration into host chromosomes^7,8^. This biological setting, in which complete, end-to-end conversion of an RNA genome is essential for viral replication, framed RTs as highly processive enzymes that operate in a continuous, template-directed manner, a view that was reinforced by subsequent studies of retrotransposons, in which RTs similarly drive copy-and-paste genome expansion through reverse transcription of mobile genetic elements^9,10^. In biotechnology, similar properties are harnessed for cDNA synthesis, where accurate and efficient copying of RNA templates is essential, motivating extensive engineering of RT variants with improved processivity, thermostability, and the ability to traverse and mechanically unfold structured RNA substrates^11,12^. Together, these observations have crystallized into a prevailing dogma: RTs function as robust polymerases that faithfully copy RNA templates into DNA, with product length defined directly by template length^13^.

Yet a growing class of RTs depart from this canonical paradigm, employing mechanisms in which DNA synthesis departs from simple end-to-end template copying. One prominent example is telomerase: rather than copying an entire RNA molecule, it uses a short, internal template embedded within a larger noncoding RNA (ncRNA) to synthesize repetitive DNA sequences at chromosome ends^14–17^, a process essential for genome stability and widely implicated in aging and cancer^18–21^. This reaction depends on repeat addition processivity (RAP), in which iterative cycles of extension and template repositioning must be precisely coordinated^22–24^, requiring strict definition of template boundaries and controlled translocation relative to the active site^25,26^. A conceptually related departure is found in retrons, bacterial genetic elements comprised of a RT and an ncRNA that function in phage defense^27,28^. Rather than copying their ncRNA end-to-end, retrons reverse transcribe only a defined internal segment of the ncRNA to generate an RNA–DNA hybrid product^29^. Although retrons do not produce long tandem repeats, they similarly rely on structured ncRNAs to define discrete DNA products^29^, illustrating that RT output can be specified by RNA architecture rather than by continuous copying along the template. Together, these systems point to a broader principle, in which RT activity need not be dictated solely by the linear extent of an RNA template, but can instead be governed by structural features of the ribonucleoprotein (RNP) complex.

This principle appears to be far more widespread in bacteria than previously appreciated, particularly within phage defense systems. Recent studies have uncovered a broad clade of non-mobile reverse transcriptases, referred to as the “Unknown Group” (UG), that are pervasive across bacterial genomes and provide antiviral immunity, leading them to be termed defense-associated reverse transcriptases (DRTs)^1,2,30,31^. The UG clade comprises three major classes with distinct genetic organizations and biochemical strategies^1,30,31^, of which Class 2 systems are distinguished by their dependence on structured ncRNAs encoded *in cis* that actively direct DNA synthesis. Across multiple Class 2 systems we have recently investigated, including DRT2, DRT3, DRT9, and DRT10, reverse transcription proceeds through iterative cycles of extension, strand separation, and template reuse, generating tandem-repeat cDNA products from short, internally embedded template regions^3–6^. This mode of synthesis bears a functional resemblance and evolutionary connection to telomerase^3,32,33^, and is exemplified by DRT10, whose template is organized around a conserved A–B–A′ pattern that directs highly periodic DNA repeat synthesis^3^. The resulting repetitive cDNA products are proposed to activate an accessory SLATT effector encoded in the same operon, coupling repeat synthesis to immune function^3^. Yet how the RNA scaffold orchestrates precise boundary definition and sustains repeated cycles of DNA synthesis remains to be explained.

Here, we define the structural and mechanistic basis of tandem-repeat DNA synthesis by DRT10. Using cryogenic electron microscopy (cryo-EM), we visualized two evolutionarily distinct RT–ncRNA complexes and uncovered an unexpected 2:1 RT–ncRNA architecture with pseudo-symmetric organization. Biochemical analyses reveal that template boundaries and repeat length are defined not by primary sequence, but by steric and geometric constraints imposed by the three-dimensional (3D) arrangement of the ribonucleoprotein complex. Finally, we show that DRT10 RT can synthesize the products of other Class 2 RT systems when provided with their corresponding ncRNA templates. Together, our findings establish a structural and mechanistic framework for RNA-templated, tandem-repeat cDNA synthesis across Class 2 RT systems, and support a broader view in which RT outputs are encoded by higher-order RNA–protein architecture.

## RESULTS

### 2:1 stoichiometry of the DRT10 RT–ncRNA complex

To understand the molecular basis of tandem-repeat cDNA synthesis by the DRT10 RT–ncRNA complex, we selected two diverse systems for structural exploration (**Fig. 1a**). Focusing first on *Eco*_1_DRT10, we reconstituted the RNP complex *in vitro* by incubating purified RT with full-length ncRNA, confirmed complex formation by analytical size-exclusion chromatography (**Fig. S1a**), and then determined its structure by cryo-EM. Preliminary micrograph analysis revealed two-dimensional (2D) class averages that exhibited apparent two-fold symmetry about a central axis (**Fig. S1b,c**), and subsequent data processing and refinement yielded a reconstruction with an overall resolution of 3.2 Å (**Fig. S1d-h**). To our surprise, analysis of the final cryo-EM map and its derived, stereochemically refined model revealed an unanticipated architecture comprising two copies of the RT enzyme bound to a single molecule of ncRNA (**Fig. 1b,c**). Closer inspection of the ncRNA immediately revealed the pseudo-symmetric nature of its secondary structure, in which three closely-positioned stem-loops (SL1–SL3 and SL4–SL6) occupy opposite sides of a central axis defined by two similarly sized loop regions, which we term Loop A and Loop B (**Fig. 1d and Fig. S2**). Loop B encompasses the template sequence and essential A–B–A′ pattern we previously linked to repetitive cDNA synthesis^3^, leaving unanswered whether Loop A is also reverse transcribed (see below).

**Fig. 1.**
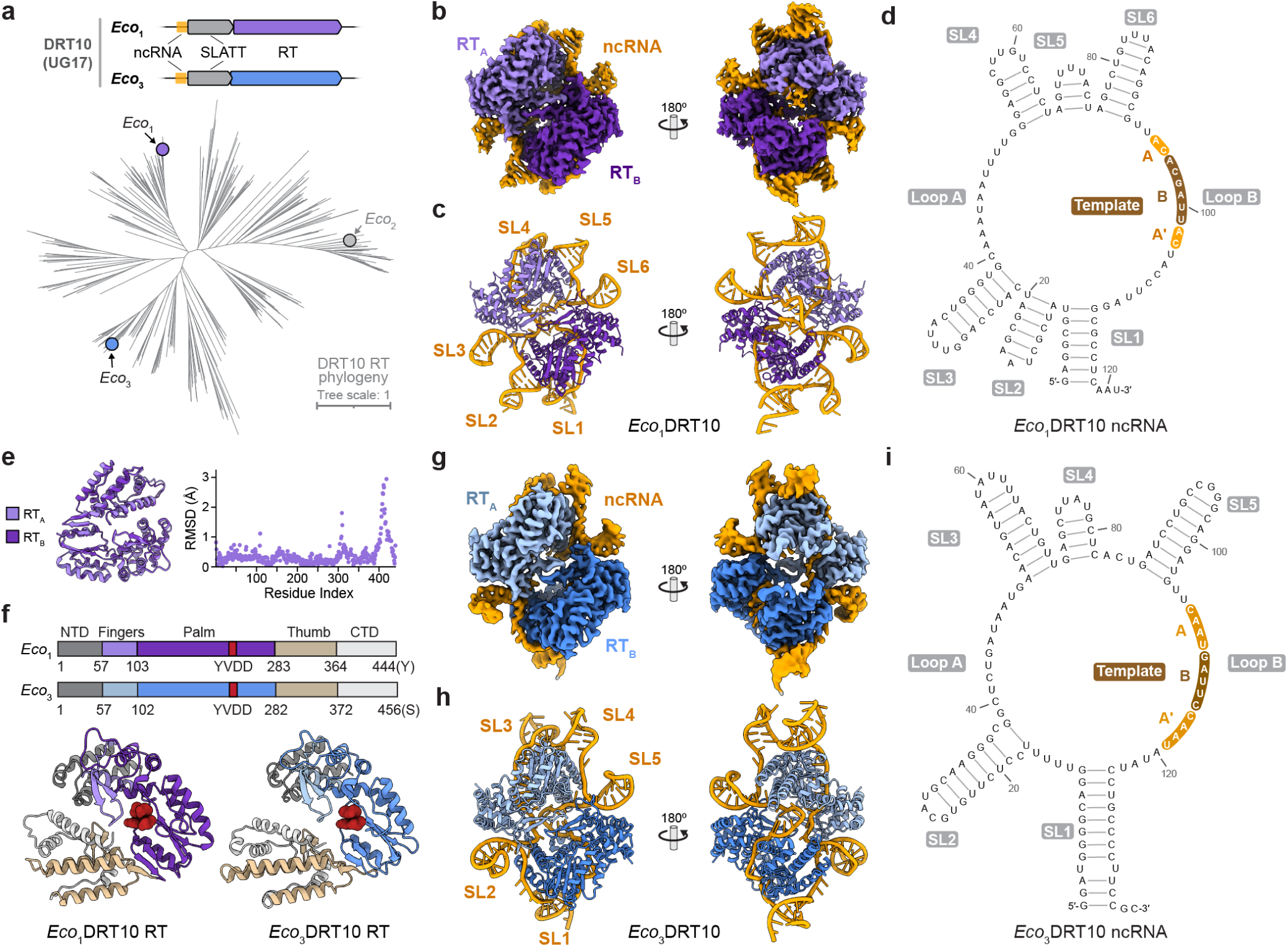
| Overall architecture and cryo-EM structures of two DRT10 RT–ncRNA complexes. **a,** Phylogenetic analysis of DRT0 homologs highlighting the divergence between *Eco*_1_ and *Eco*_3_ proteins, with operon organization shown at top. **b,** Cryo-EM reconstruction of the *Eco*_1_DRT10 RT–ncRNA complex, shown in two orientations. The two RT monomers and ncRNA are labeled. **c,** Atomic model of *Eco*_1_ RT–ncRNA complex, shown as in **b. d,** Secondary structure of the *Eco*_1_ ncRNA, with the template region, A–B–A′ pattern, and relevant features labeled. **e,** Superposition of the two *Eco*_1_RT protomers (*left*), with RMSD calculated over Cα atoms across 439 aligned residues (*right*). **f,** *Top*, RT domain organization, with residue numbering for *Eco*_1_ and *Eco*_3_ shown. *Bottom*, structures of an *Eco*_1_RT and *Eco*_3_RT monomer colored according to domain organization. **g,** Cryo-EM reconstruction of the *Eco*_3_DRT10 RT–ncRNA complex, shown in two orientations. The two RT monomers and ncRNA are labeled. **h,** Atomic model of *Eco*_3_ RT–ncRNA complex, shown as in **g. i,** Secondary structure of the *Eco*_3_ ncRNA, with the template region, A–B–A′ pattern, and relevant features labeled.

Strikingly, the two RT protomers are structurally indistinguishable, with a root-mean-square deviation (RMSD) of 0.54 Å (**Fig. 1e**), arguing against a distinct conformational state selectively activating only one monomer. Each RT adopts a canonical right-hand conformation comprising fingers, palm, and thumb domains^34^, along with a C-terminal domain (CTD) extension that harbors the priming residue (**Fig. 1f and Fig. S3a**). The ncRNA adopts a complex, figure-eight-like architecture where the lower ‘circle’ harbors SL1 (formed by pairing of the 5′ and 3′ ends), SL2, SL3, and loop A; the upper half of the figure-eight shape harbors loop B and SL4–SL6 positioned in a three-dimensional stance opposite to that of the lower half, presenting an approximately symmetric configuration overall, with a dyad axis passing through the middle of the figure eight (**Fig. 1d and Fig. S2**). With this unusual configuration, the ncRNA plays a fundamental role in coordinating the relative position of each RT, since their minimal, protein-only surface area likely provides insufficient binding energy to promote RNA-independent dimerization (**Fig. 1c and Fig. S2**). Altogether, the DRT10 ncRNA component plays a dual role as template for DNA synthesis, and as a structural scaffold to promote and maintain a dimeric RT configuration.

The *Eco*_1_DRT10 immune system provides potent antiphage defense^3^ and exhibits robust tandem-repeat cDNA synthesis *in vitro* (**Fig. S4a**), but our DNA high-throughput sequencing (HTS) analyses revealed frequent, aberrant tracts of poly-(dTdG) repeats that interrupt precise copies of the consensus 5′-AATCGTGT-3′ cDNA motif (**Fig. S4b**), as a consequence of a cryptic internal reannealing motif we described previously^3^. However, minimal mutations to remove the cryptic repeat, while mildly improving the precision of repeat addition, still yielded RAP activity far below that of the *Eco*_3_DRT10 system (**Fig. S4a,c**). Template-swapping experiments confirmed that this difference was not solely attributable to template sequence: introducing the *Eco*_1_ template into the *Eco*_3_ ncRNA increased RAP activity, whereas placing the *Eco*_3_ template into the *Eco*_1_ ncRNA significantly reduced RAP accuracy (**Fig. S4d**), indicating that these differences reflect intrinsic properties of the RT–ncRNA ribonucleoprotein itself. We reasoned that the reduced precision of *Eco*_1_ RAP would complicate a detailed study of the reverse transcription reaction mechanism, thereby motivating us to instead focus on the evolutionarily diverse *Eco*_3_DRT10 system.

We reconstituted the biochemically active *Eco*_3_ RT–ncRNA co-complex, which synthesizes precise copies of a 5′-GAATCATTG-3′ cDNA motif (**Fig. 1a and Fig. S4a**), and determined its structure by cryo-EM to an overall resolution of 3.3 Å (**Fig. 1g,h, Fig. S5a–c**). The structure adopts a remarkably similar 2:1 architecture as the *Eco*_1_ system (**Fig. 1g,h**), despite employing a ncRNA that exhibits lesser apparent symmetry in the relative positioning of its five stem-loop elements (**Fig. 1i**). As with *Eco*_1_, we observed both *Eco*_3_ RT monomers occupying nearly indistinguishable conformations with equivalent domain organization, with a RMSD of 0.72 Å over 448 aligned Cα atoms (**Fig. S5e**), and the A and B loops were again positioned in similar orientations relative to the active site within both RT monomers.

Collectively, these structures reveal an unprecedented topology for the relative coordination of two RT enzymes by a single ncRNA scaffold, and define the spatial organization of the template region within the catalytic active site. At the same time, this architecture raised a series of fundamental mechanistic questions: whether one or both RT monomers engage in productive reverse transcription, how the priming site is chosen, how the template boundaries are precisely defined, and how the system achieves rapid and processive synthesis of kilobase-length DNA repeats. In the following sections, we systematically explored these questions through a combination of biochemical, structural, and sequencing-based experiments.

### Functional asymmetry within a pseudo-symmetric RT–ncRNA complex

We developed a rapid and efficient experimental workflow to profile the effects of ncRNA perturbations on DRT10-mediated DNA synthesis. After optimizing an *in vitro* transcription and RNA cleanup protocol to produce high-quality ncRNAs from synthetic DNA fragments in microplate format, we introduced rationally designed mutations and tested their effects on reverse transcription by analyzing synthesis products via high-throughput sequencing (**Fig. S6a,b, Methods**). Of note, this method has the distinct advantage of reporting on both the accuracy and processivity of tandem-repeat cDNA synthesis, but it cannot reliably detect short abortive products, given the constraints of efficient DNA capture during library preparation. To sensitively probe these species, we alternatively used radioactive nucleotide incorporation assays (**Fig. S6b, Methods**).

We first explored the pseudo-symmetric nature of the ncRNA. Both RT monomers within the *Eco*_3_ complex contact numerous equivalent nucleotides (**Fig. 2a**), and a bioinformatic survey of predicted DRT10-encoded ncRNAs revealed broad conservation of this ncRNA architecture across the family (**Fig. 2b**), consistent with a pseudo-symmetric organization. Nevertheless, the *Eco*_3_ ncRNA harbors an uneven five stem-loops, with the location between SL1 and SL2 — equivalent to SL4 situated between SL3 and SL5 — substituted with a short 4-nt loop (**Fig. 2b**). To explore the functional importance of this asymmetry, we inserted a second copy of SL4 in this location, or removed SL4 altogether from its original location. Both variants exhibited near-WT levels of cDNA synthesis (**Fig. 2c**), in agreement with the limited interactions that occur between RT monomer A and the SL4 feature (**Fig. 2a**). SL2 and SL5 could also be swapped, consistent with their symmetric interactions with RT monomers, but we found that exchange of SL1 and SL3 led to a complete loss of tandem-repeat cDNA synthesis (**Fig. 2c**). Interestingly, the GC content differs drastically between SL1 (75%) and SL3 (25%), suggesting that increased SL1 stability might be critical for the overall integrity of the ncRNA fold.

**Fig. 2.**
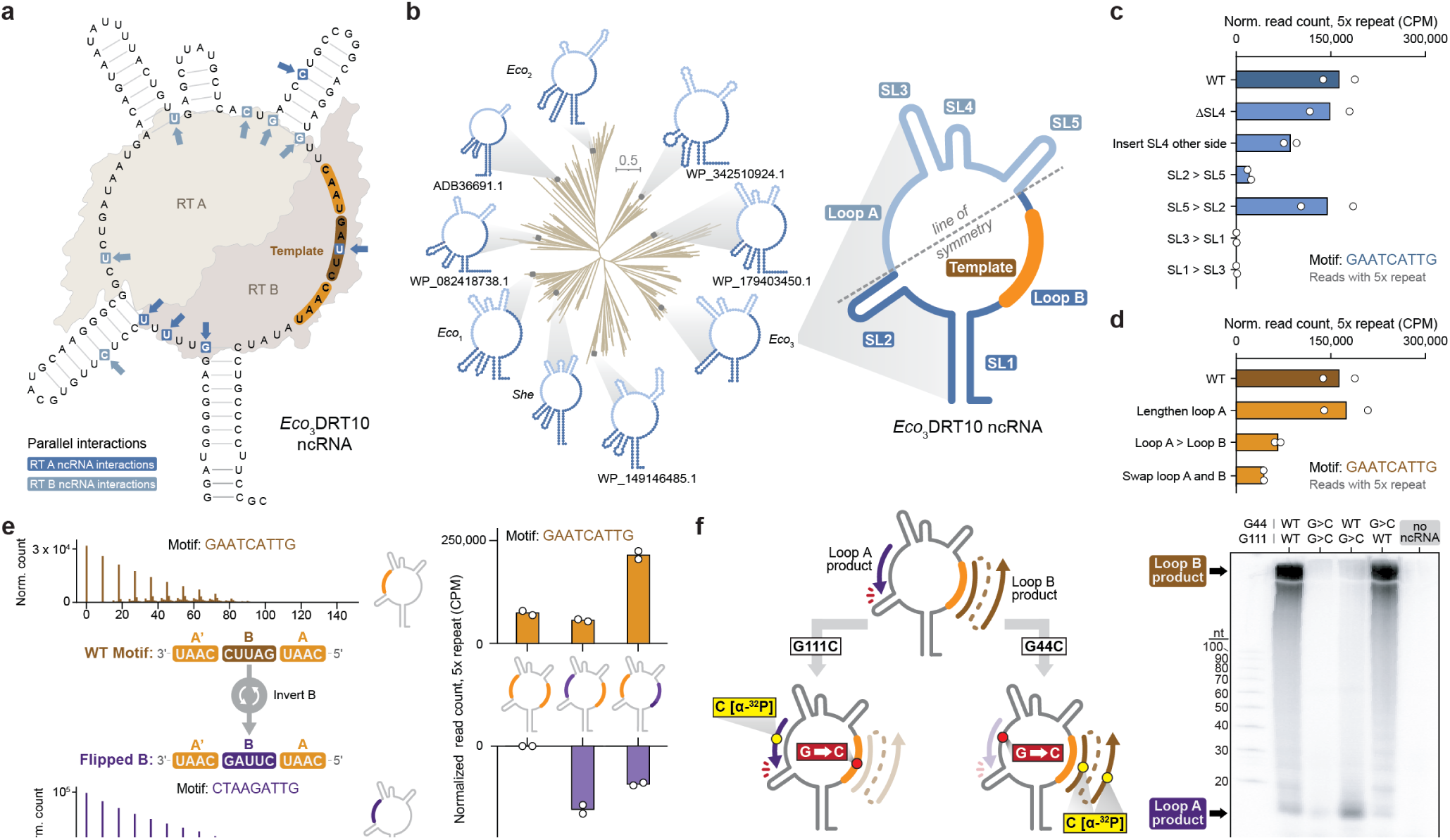
| Pseudo-symmetric architecture of the DRT10 RT–ncRNA complex. **a,** Cartoon schematic of the *Eco*_3_DRT10 RT–ncRNA complex highlighting the putative line of symmetry and symmetric RT–ncRNA interactions on opposite sides of the ncRNA. **b,** *Left*, bioinformatic analysis of ncRNA structural symmetry across diverse DRT10 systems, highlighting the conservation of the pseudo-symmetric loop and stem-loop architecture. *Right*, magnified view of the *Eco*_3_ ncRNA, with the structured stem-loops and single-stranded loop region labeled and the template located in Loop B. **c,** *In vitro* sequencing results of *Eco*_3_ ncRNA stem-loop swapping experiments. 5× repeat reads are defined as reads containing five tandem, uninterrupted copies of the consensus motif, counted and normalized to spike-in–aligned reads as counts per million (CPM). Variants include SL4 insertion and removal and SL2↔SL5 and SL1↔SL3 swaps. **d,** *In vitro* sequencing results of *Eco*_3_ ncRNA loop variation experiments. Variants include Loop A length extension, Loop A converted to Loop B, and swapping Loop A and B such that Loop B no longer contains the template. **e,** *Left*, motif position graphs for the WT and Flipped B motifs in *Eco*_3_DRT10 Loop A, in which the first occurrence of the consensus motif in each read sets the anchor position, and every subsequent occurrence of that motif is tabulated at its corresponding downstream position. The cartoons schematize inversion of the B sequence to create a unique, sequencing-distinguishable template (Flipped B). *Right*, sequencing-based quantification (5× repeat reads) for *Eco*_3_ ncRNA variants in which both loops contain a template sequence, either both WT or encoding both the WT and Flipped-B motifs in both relative orientations. **f,** *Left*, cartoon detailing the experimental setup: each loop template contains a single guanine (G44 in Loop A, G111 in Loop B), so [α-³²P]-dCTP is incorporated opposite that base, and radioactive signal therefore reports synthesis from whichever loop retains its guanine. *Right*, radiolabeled ([α-³²P]-dCTP) denaturing PAGE analysis of ncRNA variants harboring G-to-C mutations at G44 (Loop A), G111 (Loop B), or both positions. A ∼10 nt product corresponding to short, non-repetitive cDNA synthesis from Loop A is detected in the G111C single mutant. Representative data shown for experiment was repeated twice with similar results. For **c, d, e**, data are mean of *n* = 2 independent replicates.

Next, we turned our attention to Loop A (12-nt) and Loop B (19-nt) sequences (**Fig. 2d**). Prior sequencing of *in vivo* DRT10 products revealed that Loop B encodes a conserved A–B–A′ pattern that templates iterative rounds of cDNA synthesis^3^, but the dimeric RT architecture suggested that either loop, or both loops, might function as a template for reverse transcription. Increasing the length of Loop A to match Loop B had no effect on the yield of cDNA synthesis products, and copying the exact same A–B–A′ pattern in Loop A also exerted a marginal effect (**Fig. 2d**), suggesting that asymmetry in the loop sequence and/or length is not critical for regulating the enzymatic activity of the dimeric RT complex. Importantly, we were even able to completely exchange the Loop A and B sequences and nonetheless still detect robust levels of tandem-repeat cDNA products (**Fig. 2d**), proving that either RT monomer within the RT–ncRNA complex is capable of reverse transcription. We next asked whether this functional interchangeability of the two template loops was preserved across DRT10 orthologs. The *Eco*_1_ complex displays an analogous pattern of symmetric RT–ncRNA contacts (**Fig. S6c**), and exchanging *Eco*_1_ Loop A and Loop B likewise preserved high levels of cDNA synthesis (**Fig. S6d**).

We wondered whether engineered ncRNAs could template the synthesis of distinct tandem-repeat cDNA products if both Loop A and Loop B were modified to encode A–B–A′ patterns competent for repeat addition processivity. We tested this hypothesis by taking advantage of a neutral ‘watermark’ mutation in the B motif (Flipped B) that enabled differentiation of cDNA motifs arising from the WT or mutant Loop sequence (**Fig. 2e**). After first confirming that the watermarked A–B*–A′ pattern yielded near-WT levels of tandem-repeat cDNAs when inserted into Loop A (**Fig. 2e**), we tested ncRNAs encompassing functional A–B–A′ patterns in both Loops and found that, regardless of relative orientation, these variants consistently supported concurrent synthesis of both product types (**Fig. 2e**). This observation raised the possibility that intermediate DNA synthesis products might oscillate between reannealing with the A′ motif of either Loop, given their identical sequence, thereby leading to hybrid species harboring cDNA motifs templated by both B and B*. We carefully analyzed our HTS data for the existence of such chimeric products but found that levels were indistinguishable from the levels of artifactual PCR recombination generated during negative control experiments (**Fig. S6e**). Similarly, incubation of the RT enzyme with two distinct ncRNA variants containing identical template regions in Loop B, except for the B/B* motif, showed an absence of chimeric reads above background (**Fig. S6e**), further indicating that intermolecular template jumping events are rare, or nonexistent. Collectively, these results argue that tandem-repeat cDNA synthesis proceeds through repeated engagement of a single A–B–A′ template within individual RT–ncRNA complexes, with each loop-specific product generated independently of the other.

Given these results, we hypothesized that the WT Loop A sequence is likely reverse transcribed constitutively, despite lacking an A–B–A′ pattern, leading to abortive products that would be non-repetitive and too short to be captured in sequencing-based analyses. We therefore designed an alternative experiment, in which product formation was monitored by denaturing urea-PAGE using radioactive [α-³²P]-dCTP. The WT Loop A and Loop B sequence each contain a single guanine base, and reactions with a WT ncRNA correspondingly generated long, kilobase-length tandem-repeat cDNA products (**Fig. 2f**). When we introduced a G44C and G111C mutations in both Loops, radioactive product formation was abolished, but critically, mutation of only G111C in Loop B yielded a strong band at ∼10-nt, corresponding to a likely abortive product templated by Loop A (**Fig. 2f**). In agreement with this interpretation, a corresponding mutation of G44C in Loop B alone restored kilobase-length product formation while depleting the presence of the ∼10-nt abortive product (**Fig. 2f**).

Taken together, these results demonstrate that both RT monomers within the pseudo-symmetric DRT10 complex are catalytically active, but are functionally specialized: one catalyzes processive, tandem-repeat cDNA synthesis from an A–B–A′-competent template loop, whereas the other yields only short, abortive reverse transcription products.

### Insights in the DRT10 RT-ncRNA priming mechanism

We next set out to define precisely where reverse transcription initiates within each active site, and to uncover geometric constraints governing the mechanism of protein priming. Attempts to trap RT–ncRNA complexes in early states of reverse transcription through the strategic combination of deoxy- and dideoxy-nucleotides were unsuccessful, and so we adopted an alternative approach in which samples were mixed with all four dNTPs directly on cryo-EM grids, immediately prior to blotting and plunge freezing. We determined the resulting structure to a global resolution of 3.0 Å (**Fig. S7**), and although the overall architecture appeared similar to the DNA-free state, we observed unequivocal density for a single deoxynucleoside triphosphate substrate in the catalytic pocket of both RT monomers (**Fig. 3a,b**). Both nucleotides are positioned appropriately to engage in conventional Watson-Crick base-pairing with corresponding residues U43 (Loop A) and C116 (Loop B) in the ncRNA, allowing us to tentatively assign the two densities as dATP and dGTP, respectively (**Fig. 3b**). This result was in excellent agreement with radioactive [α–^32^P]dNTP incorporation experiments testing each nucleotide individually, in which we found that the *Eco*_3_RT was most efficiently labeled with dATP and dGTP, but not dTTP or dCTP (**Fig. 3c**). These observations suggest that both RT active sites are pre-organized for nucleotide incorporation, with a preference for purine nucleotides, prompting the question of how the proteinaceous nucleophile is positioned within each catalytic pocket.

**Fig. 3.**
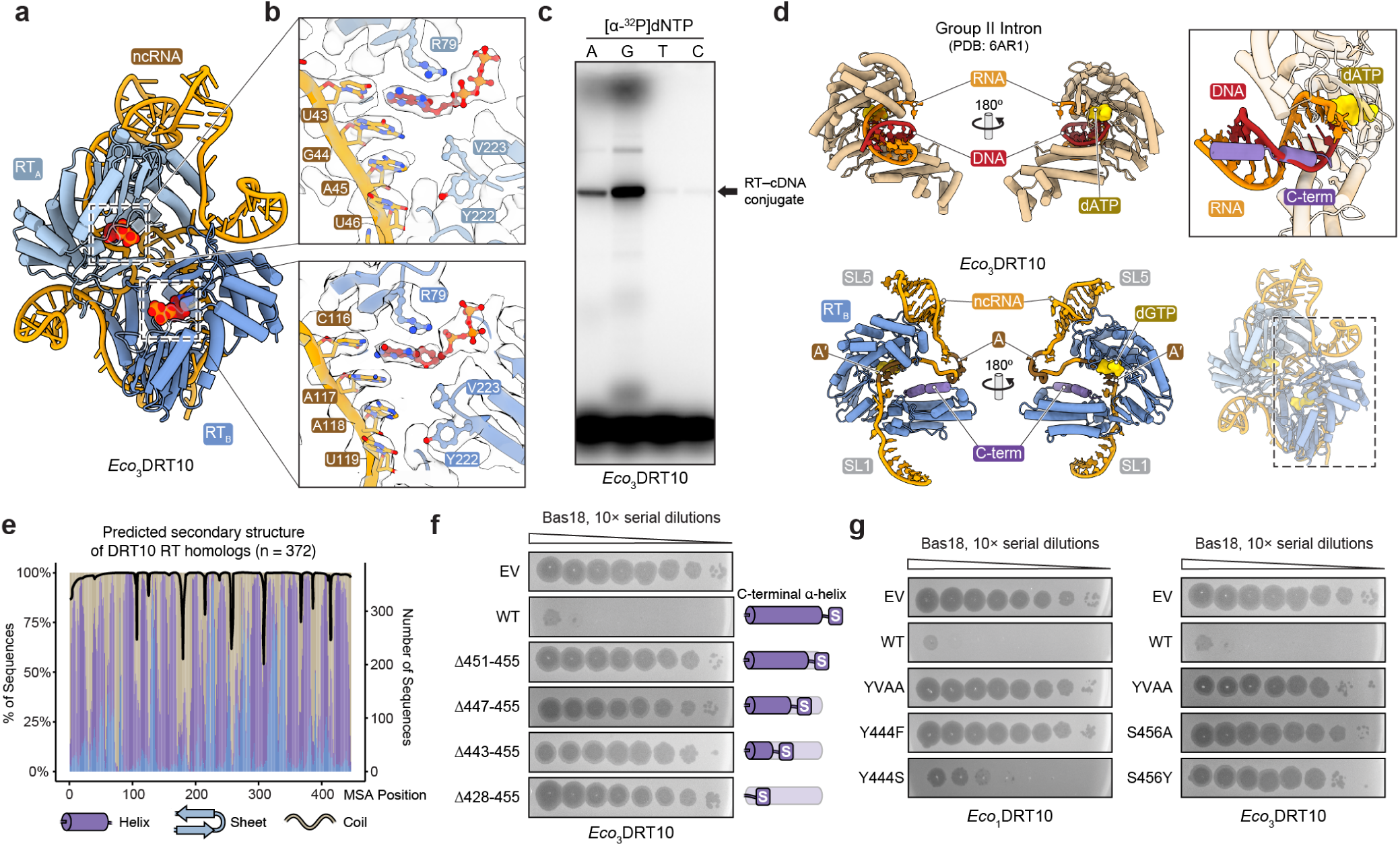
| Structure of the dNTP-bound *Eco*_3_DRT10 RT-ncRNA complex. **a,** Atomic model of the dNTP-bound *Eco*_3_DRT10 RT-ncRNA complex, with RT monomers colored blue and ncRNA colored yellow. The RT_A_-bound dATP and RT_B_-bound dGTP are highlighted as sphere representations. **b**, Close-up views of the active sites in both RT monomers. The atomic model is shown fitted into semi-transparent electron density. Density not observed in the apo structure is present adjacent to ncRNA nucleotides U34 (top) and C116 (bottom), and is assigned as dATP and dGTP, respectively, based on the shape and positioning of the density. **c**, Denaturing 12% SDS-PAGE Gel analysis of *Eco*_3_DRT10 RT binding to corresponding [α-³²P]-dNTPs. **d**, Structural comparison of the C-terminal α-helix in dNTP-bound *Eco*_3_DRT10 with the RNA–DNA duplex track of a Group II intron RT (PDB: 6AR1). Superposition of the two RTs reveals that the *Eco*_3_DRT10 C-terminal α-helix (purple) overlaps with the DNA product track of the Group II intron RT (red) (upper right panel). **e**, Predicted secondary structure across 372 DRT10 RT homologs, with emphasis on the C-terminal α-helix. **f**, Schematic of C-terminal truncation constructs of *Eco*_3_DRT10 RT, designed to truncate the α-helix while retaining the priming residue S456, with corresponding plaque assays showing loss of defense activity upon truncation. **g**, Plaque assays of priming residue swap variants between *Eco*_1_DRT10 and *Eco*_3_DRT10, showing partial loss of defense activity in the *Eco*_1_DRT10 and complete loss of defense activity in the *Eco*_3_DRT10. Representative data shown for **c, f, g,** was repeated twice with similar results.

Like DRT9-encoded RTs^5,35^, DRT10 covalently attaches DNA directly to the RT enzyme at a C-terminal Ser or Tyr residue, depending on homolog, leading to the radioactive protein band detected by SDS-PAGE analysis (**Fig. 3c**). However, unambiguous cryoEM densities could not be assigned to priming residues Y444 and S456 in our reconstructed maps for *Eco*_1_ and *Eco*_3_, respectively, suggesting an exacerbated mobility at the extreme C-termini. Clear density was visible for a highly conserved α-helix positioned immediately upstream of the C-terminal priming residue, (**Fig. 3d,e**), and remarkably, comparison with product-bound group II intron RT structures revealed that this helix sterically occupies the exact trajectory of the nascent cDNA, extending its C-terminal end directly towards the catalytic pocket (**Fig. 3d**). Together, these observations support a model in which the dNTP-bound *Eco*_3_ structure represents a pre-catalytic state, with the C-terminal helix acting as a positioning element to initiate protein-primed DNA synthesis.

To directly test the importance of priming geometry, we first explored the sensitivity of cDNA synthesis to the positioning of the priming residue relative to the C-terminus. We generated a series of *Eco*_3_ RT C-terminal helix truncations while preserving a terminal serine residue, and found that even a minimal 5-amino acid deletion was sufficient to completely abolish phage defense (**Fig. 3f**). We next examined whether the chemical identity of the priming residue itself imposes similar constraints, given that distinct DRT10 homologs use either serine or tyrosine. Interestingly, substitution of the *Eco*_3_ priming serine with tyrosine (S456Y) resulted in a complete loss of phage defense activity, whereas the reciprocal mutation in *Eco*_1_ (Y444S) largely retained defense activity (**Fig. 3g**). This observation reveals a surprising degree of chemical flexibility in the priming mechanism, suggesting that the RT active site can accommodate distinct, hydroxyl-containing nucleophilic residues within certain structural contexts.

Together, these findings establish that placement of the priming residue at the C-terminus is a strict requirement for protein-primed initiation in DRT10, while its chemical identity is more flexible. With initiation defined, we next turned to the question of how the active site enforces precise template usage across iterative cycles of extension and resetting, to generate uniform tandem repeats during RAP.

### Geometric definition of template boundaries during tandem-repeat synthesis

A defining feature of DRT10 is its ability to repeatedly synthesize the same cDNA motif from a short, internally embedded segment of the ncRNA template, generating tandem-repeat products with remarkably uniformity. This behavior imposes a stringent mechanistic requirement: the RT must terminate synthesis at a precise position and reinitiate at the template start during each subsequent cycle, without loss of register or read-through into adjacent RNA sequences. We set out to understand how these template boundaries are established and enforced, at nucleotide resolution (**Fig. 4a**).

**Fig. 4.**
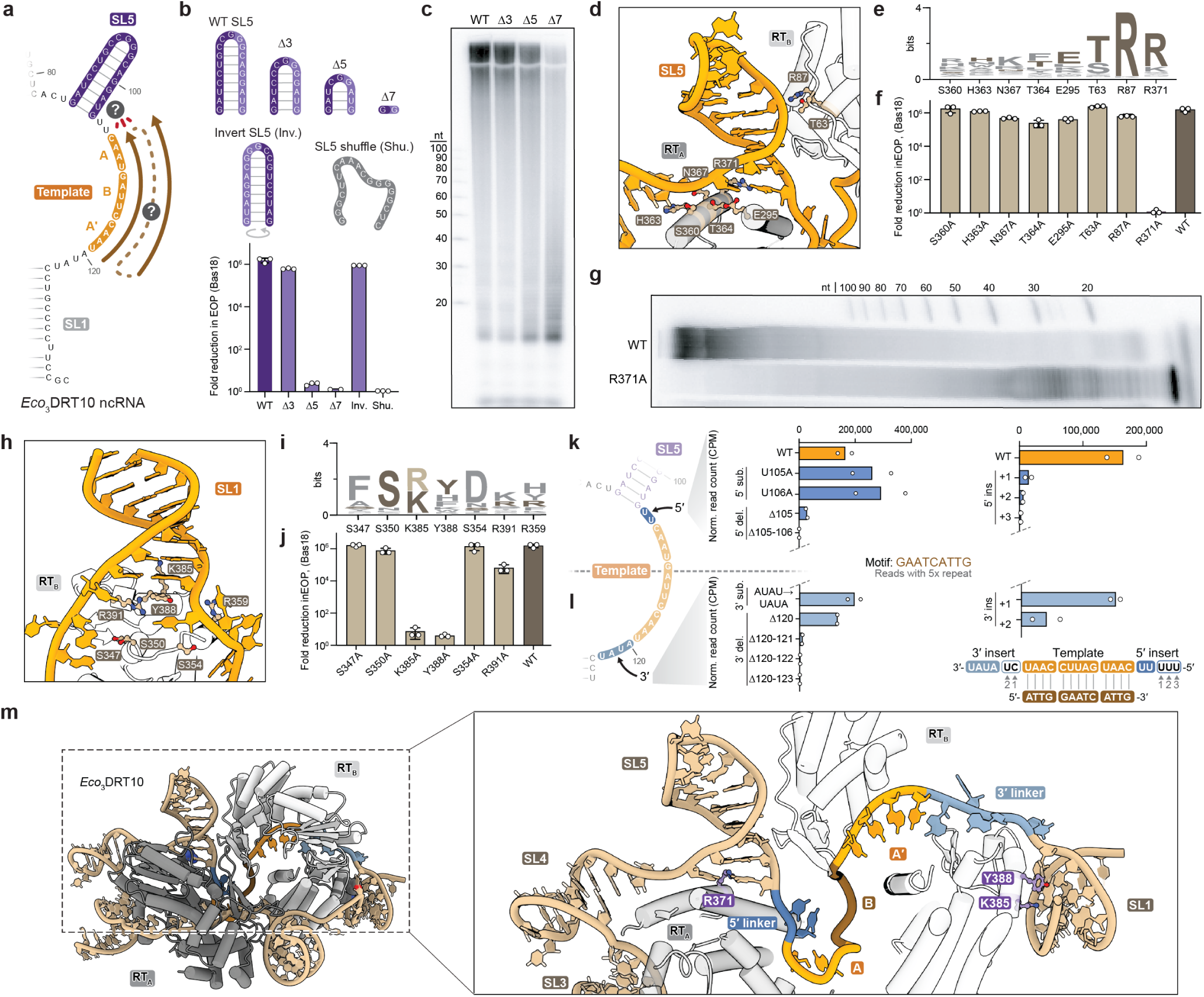
| Template boundary determinants of DRT10 repeat synthesis. **a,** Schematic illustrating the question of how the RT is halted at the 5’ template boundary and resets to the 3’ end for processive repeat addition. **b,** *Top*, cartoon schematizing the SL5 truncation, inversion, and shuffling ncRNA variants. *Bottom*, *in vivo* phage defense assays for these variants, quantified as fold reduction in efficiency of plating (EOP) relative to an EV control. **c,** Radiolabeled (α-³²P-dCTP) denaturing PAGE analysis of *in vitro* cDNA synthesis products from SL5-truncated ncRNA complexes, revealing accumulation of a short (∼13–14 nt) product consistent with read-through past the template boundary. Representative data shown for experiment was repeated twice with similar results. **d,** Diagram of RT–ncRNA interactions at the SL5 boundary element and surrounding nucleotides, with interacting residues labeled. **e,** Sequence conservation (weblogo) of RT residues that contact SL5 across diverse DRT10 homologs. **f,** *In vivo* phage defense assays for alanine mutations at each of the 8 SL5-interacting RT residues, identifying R371 as the sole residue essential for defense. **g,** Radiolabeled ([α-³²P]-dCTP) denaturing PAGE analysis of *in vitro* cDNA synthesis by the R371A mutant, showing complete loss of full-length product and strong accumulation of a ∼10 nt species. Representative data shown for experiment was repeated twice with similar results. **h,** Diagram of RT–ncRNA interactions at the SL1 boundary element and surrounding nucleotides, with interacting residues labeled. **i,** Sequence conservation (weblogo) of RT residues that contact SL1 across diverse DRT10 homologs. **j,** *In vivo* phage defense assays for alanine mutations at SL1-interacting RT residues, identifying two residues essential for defense. **k,** Schematic of the 5′ linker (U105–U106, between SL5 and the template), with accompanying sequencing-based quantification of cDNA products (5× repeat reads) from linker sequence-identity swaps, progressive linker deletions, and insertions that lengthen the linker, as detailed in the cartoon below. **l,** Schematic of the 3′ linker (A120–U123, between the template and SL1), with accompanying sequencing-based quantification of cDNA products (5× repeat reads) from linker sequence-identity swaps, progressive linker deletions, and insertions that lengthen the linker, as detailed in the cartoon below. **m,** Structural view of the template-boundary regions, with the 5′ and 3′ linkers and their anchoring stem-loop contacts highlighted. For **b, f, j**, data are mean ± s.d. of *n* = 3 technical replicates. For **k, l,** data are mean of *n* = 2 independent replicates.

We hypothesized that SL5 serves as a structural boundary element that halts reverse transcription at residue C107, at the 5’ template edge. To assess its importance, we performed progressive base-pair truncations from the top of the stem while preserving the sequence identity of the paired region. Deletion of more than 3 bp largely abolished phage defense activity, with concomitant deleterious effects on the yield and length distribution of cDNA synthesis products (**Fig. 4b,c and Fig. S8a**). In stark contrast, inversion of the SL5 sequence, in which base-pairing was maintained but nucleotide identities were swapped, had no detectable effect on phage defense (**Fig. 4b**), indicating that the SL5 secondary structure, but not its primary sequence, is critical for tandem-repeat cDNA synthesis and phage defense. Consistent with this, complete shuffling of the SL5 sequence fully abrogated defense function (**Fig. 4b**). Closer inspection of radiolabeled gel data with SL5-truncated reactions revealed the accumulation of short products ∼15-nt in length (**Fig. 4c**), indicative of potential read-through beyond the WT template boundary up to the base of SL4, followed by a failure to accurately reset and perform additional rounds of repeat addition. High-throughput sequencing results corroborated this interpretation, demonstrating that progressive SL5 truncations substantially increased the frequency of reads containing 3’ ends templated by ncRNA nucleotides beyond the template boundary (**Fig. S8b**).

We next sought to identify key RT residues that mediate SL5 interactions and enforce this boundary element. Structural analysis revealed a network of contacts between the RT and the base of SL5 and adjacent nucleotides (**Fig. 4d**). To assess their functional importance, we performed alanine scanning mutagenesis of eight candidate residues and evaluated their effects on phage defense. Activity was abolished with only a single mutation of the highly conserved R371 (**Fig. 4e,f**) that engages the SL5 base through bidentate hydrogen bonds with G104, which itself forms a non-canonical Hoogsteen conformation with G86, imparting a distinctive local geometry (**Fig. S8c**). Consistent with this model, biochemical characterization of a R371A mutant revealed a complete loss of kilobase-length cDNA synthesis accompanied by accumulation of a short (∼10-nt) product (**Fig. 4g**), suggesting that the SL5 anchoring interaction is required for accurate template resetting and processive repeat synthesis. Interestingly, despite strong conservation of the G–G Hoogsteen pair at this position (90–95% by covariance analysis), our mutational analysis revealed minimal dependence on nucleotide identity, as substitution with alternative base combinations, including a putative A–G Hoogsteen pair, exerted minimal effects on cDNA synthesis (**Fig. S8d**). These results support a model in which a conserved RT–RNA anchoring interaction enforces the template boundary at the 5’ end of the template.

We next asked whether analogous RT–RNA interactions with SL1 also help to define the 3’ boundary of the template. Structural analysis revealed a network of seven key interactions within this region (**Fig. 4h,i**), and genetic perturbation experiments highlighted two amino acid residues (K385 and Y388) whose substitution led to a loss of defense (**Fig. 4j**). These results suggest that like SL5, SL1 engages the RT through a critical set of interactions that are important for proper template positioning and definition.

The ncRNA contains additional nucleotides immediately adjacent to both SL1 and SL5 that are not reverse transcribed into cDNA, raising a key mechanistic question: what determines the precise spacing between these anchoring elements and the nucleotides defined as part of the template? To address this, we examined the single-stranded RNA segments that flank the template region, namely U105–U106 (the 5′ linker) and A120–U123 (the 3′ linker). Mutation of either linker exhibited a negligible effect on tandem-repeat cDNA synthesis, but the deletion of two or more nucleotides from either the 5′ or 3′ linker abolished product formation (**Fig. 4k**). Interestingly, a single 5′ deletion (ΔU105) caused a severe 83.5% decrease in detectable tandem-repeat products *in vitro* (**Fig. 4k**), but had a minimal effect on phage defense activity *in vivo* (**Fig. S8e**), suggesting a non-linear relationship between DNA synthesis activity and the thresholding required to activate the SLATT effector protein for immunity. Even more intriguingly, the 2-nt deletion eliminated detectable WT motif products and abolished phage defense but nevertheless appeared to retain high synthetic yields of kilobase-length DNA, as visualized by radioactive urea-PAGE experiments (**Fig. 4k and Fig. S8e,f**). Closer inspection of the raw sequencing data revealed that this 5′ linker deletion (ΔU105–U106) produces an alternative repeat unit, CAT, appearing as CATCATCAT in the weblogo (**Fig. S8g**). The 5′ linker deletion products also displayed an alternative A–B–A′ pattern consistent with the rescaled template (**Fig. S8g**), with synthesis now terminating at the same distance from SL5 as in the WT ncRNA, set by the anchoring interaction with the stem-loop. However, this fixed spacing places the 5′ boundary 2 nt closer to the 5′ edge of the original template, yielding a 3-nt repeat instead of a 9-nt repeat **(Fig. S8g)**. Together, these results suggest a remarkably flexible system for template boundary definition, in which a geometrically constrained window, measured from the SL5 anchor rather than a fixed sequence, defines repeat synthesis.

To further explore this feature, we performed the reciprocal experiment by lengthening the 5′ linker. Consistent with a strict requirement for precise boundary position, insertion of one or more uridine residues led to a rapid loss of tandem-repeat cDNA synthesis products (**Fig. 4l**). Analysis at the level of individual repeat units revealed a distinct underlying defect. By extracting single motifs from sequencing reads and examining the nucleotide immediately 3′ of the WT motif, we observed frequent read-through events, characterized by the addition of adenosine residues templated by uridine (**Fig. S8h**). These extensions effectively shift the template boundary but do not generate a compatible A–B–A′ architecture required for iterative template resetting. As a result, nucleotide incorporation can proceed beyond the canonical boundary, but the misregistered products fail to support effective repeat addition processivity and instead collapse into short, non-productive DNA fragments. Together, these findings indicate that even subtle boundary shifts disrupt the ability of the RT to re-engage the template, aborting the repeat synthesis cycle.

We next applied the same set of perturbations to the 3′ linker (A120–U123), asking whether the same strict positional requirements hold true at the opposing boundary. The single deletion ΔA120 had little effect on tandem-repeat cDNA synthesis, in contrast to the substantial loss observed for the equivalent ΔU105 at the 5′ linker (**Fig. 4k**). Insertions in the 3′ linker were also accommodated to a notable degree: a 1-nt insertion yielded near-WT product levels, while a 2-nt insertion still produced detectable tandem-repeat cDNAs (**Fig. 4l**). We interpret this relative flexibility to reflect an additional layer of boundary definition at the 3′ side, in which the A/A′ microhomology between the template and the nascent cDNA helps to register the position from which each new repeat initiates, partially decoupling repeat synthesis from strict 3′ linker length. Nevertheless, larger 3′ deletions (Δ120–121 and beyond) abolished product formation (**Fig. 4k**), indicating that steric constraints between SL1 and the template still impose a minimum linker length compatible with productive repeat synthesis. Taken together, the 5′ and 3′ linker perturbation experiments reveal an asymmetric but coordinately enforced boundary architecture, in which the 5′ end is defined by a geometrically rigid SL5-anchored window and the 3′ end is subject to partial sequence-level correction, yet both boundaries must fall within a narrow positional tolerance to support processive tandem-repeat synthesis (**Fig. 4m**).

To assess whether these boundary-defining interactions are conserved across DRT10 homologs, we examined the corresponding regions in the *Eco*_1_ RT–ncRNA system. The overall organization of the SL6 (equivalent to *Eco*_3_ SL5) and SL1 boundary elements, and their positioning relative to the RT active site, are notably similar to those observed in *Eco*_3_ (**Fig. S8i**). In particular, the base of SL6 engages an analogous set of contacts with the RT, and SL1 is similarly positioned to define the opposing boundary of the template region. The presence of this architecture in two evolutionarily divergent DRT10 systems suggested that template boundaries are defined by a common set of RT–RNA interactions, but left open the question of what constraints, if any, are imposed on the sequence of the template itself.

### Conserved logic for tandem-repeat cDNA synthesis across Class 2 RT systems

Our earlier perturbations of the A–B–A′ motif suggested that DRT10 is broadly tolerant of sequence variation as long as the A–B–A′ pattern is preserved. To map the limits of this flexibility, we systematically varied B length, and GC content. Substantial changes in both parameters were compatible with robust tandem-repeat cDNA synthesis, provided the overall A–B–A′ pattern was preserved (**Fig. 5a,b**), highlighting the functional constraints on the A–B–A′ architecture. We next asked if this tolerance is reflected in the RT–ncRNA interface itself. The RT makes minimal sequence-specific contacts with the template region, and a mutational scan of the four template-interacting residues identified only one, N197, as essential for phage defense (**Fig. 5c**). Together, these results support a model in which repeat synthesis is governed not by the identity of the template sequence, but by a geometrically defined template window flanked by anchoring stem-loops, with A–A′ microhomology directing iterative template resetting between cycles of extension (**Fig. 5d**).

**Fig. 5.**
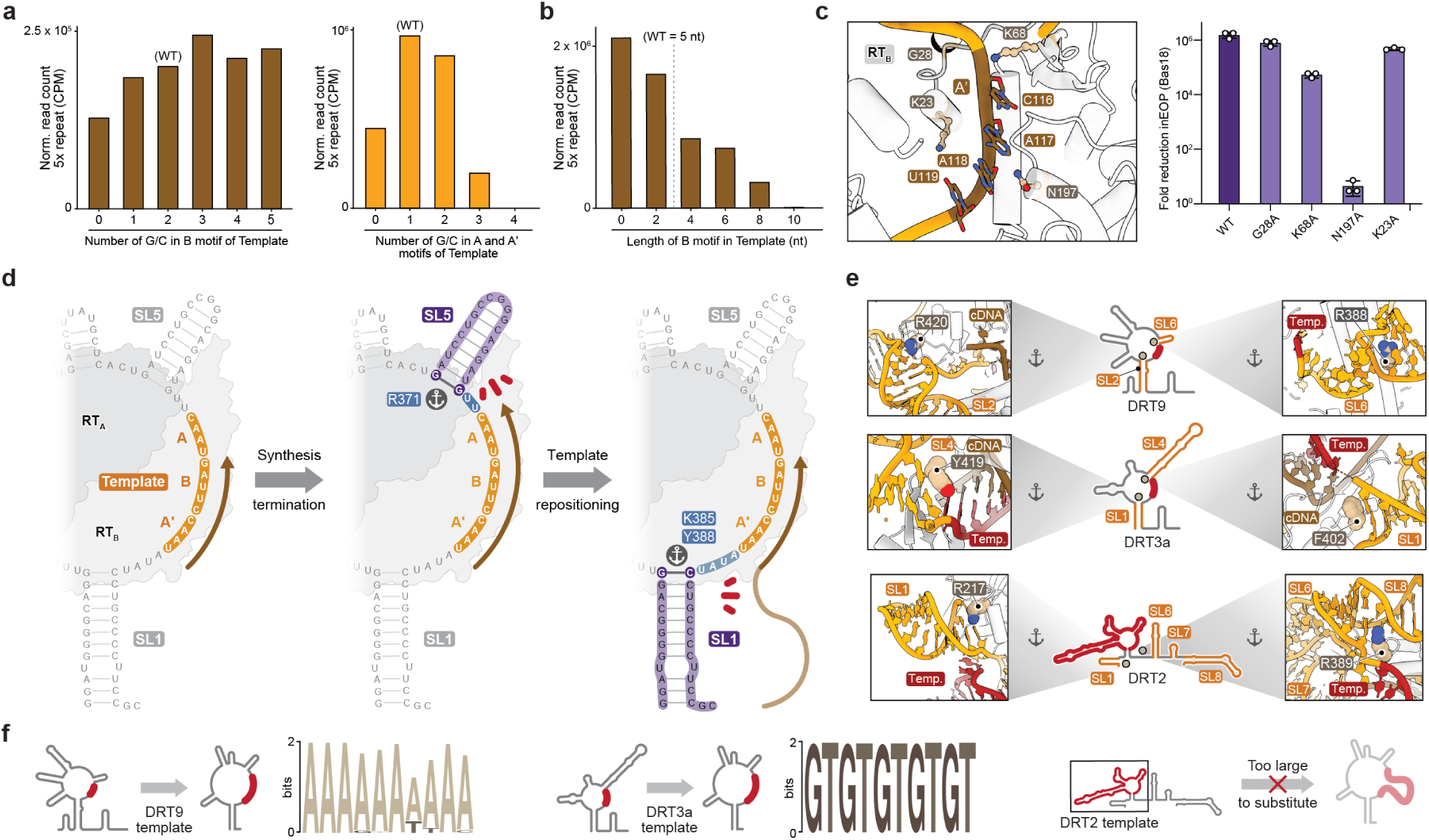
| Conserved mechanism of template boundary definition across DRT systems. **a,** Sequencing-based quantification of *in vitro* cDNA products (5x repeat reads) from ncRNA variants with progressive GC-content modulation across the loop B template (left) and across the flanking A/A′ regions (right), with the unmodulated region retained as WT in each case. Variant motifs are listed in **Table S7**. **b,** Sequencing-based quantification of *in vitro* cDNA products (5x repeat reads) from ncRNA variants with progressive shortening and extension of the B motif of the template region. Variant motifs are listed in **Table S7**. **c,** *Left*, structural view of RT residues contacting the loop B template region of the ncRNA. *Right*, *in vivo* phage defense activity of alanine substitutions at these residues, quantified as fold reduction in efficiency of plating (EOP) relative to an EV control. Data are mean ± s.d. of *n* = 3 technical replicates. **d,** Cartoon model of the proposed repeat synthesis mechanism, depicting anchoring interactions on either side of the loop B template and the flanking linker regions that permit template translocation through the RT active site. **e,** Structural views of RT–ncRNA interactions in DRT9 (PDB 9VMA), DRT3a (PDB 9Z6Z), and DRT2 (PDB 9C0I), analogous to the anchoring contacts identified in DRT10 at the stem-loop bases that flank the template region. **f,** MEME motif analysis of cDNA products from ncRNA variants in which the WT template sequence was replaced with sequences predicted to direct synthesis of cDNAs characteristic of other Class 2 RT systems. Data shown in **a, b, f,** was repeated twice with similar results.

We next asked whether this architectural logic extends to other Class 2 DRT systems. Remarkably, our inspection of recently determined RT–ncRNA structures from DRT2^36^, DRT3^37^, and DRT9^38^ revealed the same underlying arrangement: in each case, two flanking stem-loops frame a short single-stranded template region, with each stem-loop anchored at its base by RT contacts (**Fig. 5e**). At the 5′ stem-loop, a single RT residue is positioned to make this contact, R420 in DRT9 and Y419 in DRT3a, occupying the same role we defined for R371 in DRT10^37,38^. Functionally, DRT3 and DRT9 likewise generate defined tandem-repeat products, and each of their template-bearing regions can be decomposed into an A–B–A′-like configuration. This shared architecture is striking given the very different oligomeric states of these systems: DRT9 is a 6:6 RT–ncRNA complex (a dimer of trimers)^5,35,38^, DRT3 is a 6:6:6 RT–ncRNA–RT complex combining two distinct RT enzymes^37^, and DRT2 is a monomer^36^. The DRT2-encoded ncRNA comprises a substantially longer B-equivalent template region, and closer inspection of the DRT2 structure revealed a noncanonical G–G conformation at the 5′ end of this region contacted by R217 (**Fig. 5e**), directly paralleling the G–G conformation and R371 in *Eco*_3_DRT10. These structural parallels suggest that a common architectural framework underlies tandem-repeat cDNA synthesis throughout the Class 2 RT family, despite substantial divergence in protein sequence, ncRNA sequence, and oligomerization state.

If a single architectural framework truly underlies repeat synthesis across these systems, we hypothesized that the DRT10 scaffold should therefore accommodate template sequences drawn from other Class 2 RTs and direct synthesis of their characteristic repeat products. To test this, we replaced the native *Eco*_3_DRT10 template motif with sequences derived from DRT3 and DRT9. Strikingly, the *Eco*_3_ RT–ncRNA complex retained the ability to generate distinct tandem-repeat cDNA products from these chimeric substrates, including poly-dA (DRT9) and poly-(dTdG) (DRT3) (**Fig. 5f**). Together, these findings reveal a common architectural logic underlying diverse Class 2 RT systems, in which sequence-diverse templates are accommodated within a conserved framework of anchored boundary elements that enables iterative repeat addition.

## DISCUSSION

Our structural characterization of the DRT10 RT–ncRNA complex reveals an unanticipated architectural principle underlying tandem-repeat reverse transcription. Two structurally indistinguishable RT monomers are coordinated by opposite sides of a single, pseudo-symmetric ncRNA, yielding a 2:1 stoichiometry with no known parallel among other RT enzymes. The ncRNA in this configuration serves a dual role, simultaneously encoding the template and scaffolding the catalytic machinery that reads it. This organization is preserved across the evolutionarily divergent *Eco*_1_ and *Eco*_3_ systems, which differ substantially in ncRNA sequence and stem-loop arrangement, indicating that the 2:1 architecture is a fundamental requirement for DRT10 function rather than an idiosyncratic feature of one homolog.

Does the structural symmetry of the ncRNA imply functional equivalence between the two RT monomers? Our biochemical dissection reveals that both RT monomers are catalytically active, but functionally specialized. The monomer engaging Loop B generates kilobase-length tandem-repeat cDNA, while the monomer engaging Loop A produces constitutive abortive products ∼10 nt in length. Loop-swap experiments demonstrate that this asymmetry is not intrinsic to the protein but emerges from whether the template loop encodes an A–B–A′ configuration permissive for microhomology-directed repositioning. Swapping Loop A and Loop B transfers processivity to the formerly abortive active site, and when both loops are engineered to carry distinct A–B–A′ sequences, each active site synthesizes its own product independently, with no detectable chimeric products. The decision between processive and abortive synthesis is made entirely at the level of RNA sequence, with two conformationally equivalent monomers reading out different functional programs.

DNA products synthesized by DRT10 are covalently linked to the enzyme itself, and protein-primed initiation is determined by positioning of the C-terminal helix, which initially occupies the same site that is ultimately occupied by the RNA–DNA heteroduplex. The priming residue itself, a serine in *Eco*_3_ and a tyrosine in *Eco*_1_, is disordered in the cryo-EM maps, consistent with the mobility required for the transition from initiation to elongation. DRT9 similarly uses a C-terminal RT residue for protein priming^5,35^, raising the possibility that protein-primed initiation from the C-terminus is a more broadly adopted strategy within the Class 2 UG family. Eukaryotic telomerase, by contrast, which similarly uses a ncRNA to synthesize long tandem-repeat cDNAs, initiates on a pre-existing telomeric DNA substrate and relies on a TEN domain anchor site and IFD-TRAP composite channel to capture and retain that substrate during translocation^39,40^. We propose that protein-primed initiation in DRT10 may partially compensate for the absence of any necessary accessory factors, thereby ensuring that the substrate-capture problem is resolved through physical RT–DNA coupling rather than a dedicated anchor domain. The structural basis for the transition from initiation to processive elongation remains an open question, and will require capturing an elongation-state structure in which the C-terminal helix has vacated the active site.

To resolve the mechanism of tandem-repeat DNA synthesis, we used mutational perturbations to demonstrate that the physical template window is defined by the distance between two RNA structural anchors — SL5 at the 5′ boundary and SL1 at the 3′ boundary — each engaged by conserved protein contacts that tightly anchor the RT. At the 5′ boundary, R371 makes bidentate hydrogen bonds to G104 via a non-canonical Hoogsteen G–G conformation, a role functionally analogous to the conserved arginine in the *Tetrahymena* TERT RNA-binding domain that contacts the base of the TBE stem-loop^41^. Intriguingly, other structurally characterized Class 2 systems each feature an equivalent contact at the base of a template-flanking stem-loop^3,4,6,35–37^. That three diverse Class 2 systems rely on a single protein–RNA contact at this position, varying only in the identity of the residue, suggests this interaction is a structural requirement for template boundary definition in the RT family. The 3′ boundary is less precisely tied to linker length, consistent with A–A′ microhomology providing an additional positional constraint. This two-anchor, two-linker architecture is reminiscent of the RNA accordion model of telomerase, in which single-stranded RNA elements flanking the template undergo reciprocal extension and compaction during translocation^25^.

The structural details reported here extend the biochemical evidence for shared mechanisms of tandem-repeat synthesis between bacterial DRT10 and eukaryotic telomerase^3^, despite their divergent protein scaffolds. We propose that DRT10 represents a structurally minimal version of the same molecular logic, whereby processive repeat synthesis is achieved with only the RT domain and boundary-anchoring contacts at conserved stem-loops flanking the template, without requiring the TEN domain, IFD-TRAP composite channel, or C-terminal extension that elaborate TERT in eukaryotes. This view is consistent with protein domain additions representing later eukaryotic innovations layered onto an RNA architectural foundation that was already sufficient for repeat addition processivity in the bacterial ancestor.

## METHODS

### Plasmid and E. coli strain construction

All strains and plasmids used in this study are described in Supplementary Tables 1 and 2. The *Eco*_1_ and *Eco*_3_ DRT10 operons, including their native upstream and downstream flanking sequences, were chemically synthesized (GenScript) and cloned into a pACYC184 plasmid backbone by Gibson Assembly. Derivative plasmids encoding RT and ncRNA mutations were generated by around-the-horn PCR, or PCR and Gibson assembly of fragments. All plasmids were cloned and propagated in *E. coli* NEB Turbo (sSL0410) and verified by Sanger or whole-plasmid sequencing. Manual inspection of the multiple sequence alignment for closely related DRT10 homologs revealed a likely misannotation in the translation start site of *Eco*_3_ DRT10 RT (NCBI accession OKX54664), resulting in the inclusion of two extra amino acids at the N-terminus. Amino acid numbering in this manuscript has been modified to reflect this correction (i.e., S458 in the NCBI record is numbered here as S456).

### Phage propagation and plaque assays

Bacteriophages were propagated in liquid culture by picking single plaques and inoculating LB media, supplemented with 5 mM CaCl_2_ and 5 mM MgSO_4_, containing *E. coli* MG1655 cells diluted 1:100 from overnight cultures. After 4–5 hours of incubation with shaking at 37 °C, lysates were centrifuged at 4,000 × g for 10 min to pellet cell debris, and chloroform was added to a final concentration of 5% to lyse residual bacteria. Supernatants were passed through a sterile 0.22 µm filter and stored at 4 °C.

Small-drop plaque assays were performed as previously described^5^. Briefly, *E. coli* K-12 strain MG1655 (sSL0810) was transformed with the indicated plasmids, and individual colonies were used to inoculate LB media and were grown overnight at 37 °C. Cultures were mixed with molten soft agar (0.5% agar in LB media supplemented with 5 mM CaCl_2_ and 5 mM MgSO_4_) at 45 °C and poured over solid bottom agar (1.5% agar in LB media containing the appropriate antibiotic) in a Petri dish. 10× serial dilutions of phage in LB media were spotted onto the surface of the soft agar lawn, and plates were incubated at 37 °C for 8–16 hours to allow plaque formation. Plaque forming units (PFU) mL^−1^ were calculated by counting plaques at the lowest dilution showing distinct plaque formation and dividing by the volume of phage spotted, with appropriate adjustment for the dilution factor. When individual plaques were indistinguishable, a count of 50 plaques was assigned at the lowest dilution showing visible lawn clearance. Phage defense activity was assessed by calculating the fold reduction in efficiency of plating (EOP), defined as the PFU mL^−1^ obtained on a lawn of empty vector (EV)-transformed control cells divided by the PFU mL^−1^ obtained on a lawn of defense system-expressing cells.

### DRT10 RT purification

RT from *Eco*_1_ DRT10 and *Eco*_3_ DRT10 systems was purified as previously described^3^, with minor modifications. His-GST-tagged RT expression constructs (WT or mutant) were cloned and transformed into *E. coli* BL21-AI cells. Cultures were grown in 2xYT media at 37 °C to OD_600_ ∼0.6, induced with 0.25 mM IPTG, and grown overnight at 16 °C. Cells were pelleted by centrifugation at 3,000 × g for 25 min at 4 °C and lysed by sonication in Lysis Buffer (20 mM HEPES-KOH pH 8.0, 500 mM NaCl, 5 mM MgCl_2_, 5% glycerol, 10 mM imidazole, 0.1% Triton-X-100, 1 mM TCEP, 1× cOmplete EDTA-free Protease Inhibitor cocktail). Lysates were clarified by centrifugation at 10,000 × g for 30 min at 4 °C, incubated with Ni-NTA resin (QIAGEN) for 1 h at 4 °C (1 mL bead volume per 1 L initial culture), and loaded onto an Econo-Pac column (BioRad). The resin was rinsed with 10 column volumes of Wash Buffer (20 mM HEPES-KOH pH 8.0, 500 mM NaCl, 5 mM MgCl_2_, 5% glycerol, 20 mM imidazole, 1 mM TCEP) prior to elution with Elution Buffer (20 mM HEPES-KOH pH 8.0, 500 mM NaCl, 5 mM MgCl_2_, 5% glycerol, 300 mM imidazole, 0.1% Triton-X-100, 1 mM TCEP). Desired fractions were combined and subjected to TEV protease cleavage for 5 h at 4 °C, using 2.5 µg TEV protease per 100 µg target protein. Samples were diluted to 300 mM NaCl immediately following cleavage and applied to a HiTrap Heparin 5 mL column (Cytiva), eluting with a continuous gradient from Buffer A (20 mM HEPES pH 7.5, 350 mM NaCl, 1 mM TCEP, 5% glycerol) to Buffer B (20 mM HEPES pH 7.5, 1 M NaCl, 1 mM TCEP, 5% glycerol). Desired fractions were combined, concentrated using a 10 kDa MWCO Amicon Ultra Centrifugal Filter (Millipore Sigma), snap-frozen in liquid nitrogen, and stored at −80 °C. Protein purity was assessed by denaturing 10% SDS-PAGE.

### ncRNA variant generation

Double-stranded DNA fragments encoding a T7 promoter followed by the ncRNA sequence of interest (WT or variant) were ordered as synthetic gene fragments (IDT eBlocks) and arrayed in 96-well plates. Fragments were used directly as templates for *in vitro* transcription with T7 RNA polymerase. Reactions (25 µL) contained 1× TS buffer (150 mM Tris-Cl pH 8.1, 0.005% Triton X-100, 2.9 mM spermidine), 5 mM each of ATP, CTP, GTP, and UTP, 10 mM DTT, 0.002 U µL^-1^ inorganic pyrophosphatase, 14.4 mM MgCl_2_ , 100 ng template DNA, 0.1 mg mL^-1^ T7 RNA polymerase, and 1 U µL^-1^ SUPERase·In RNase inhibitor. Reactions were incubated at 37 °C for 12 h, treated with TURBO DNase for 45 min at 37 °C, and heat-inactivated by addition of EDTA to 15 mM final and incubation at 75 °C for 10 min. RNA products were purified using the Zymo RNA Clean & Concentrator-25 kit (R1017) according to the manufacturer’s instructions, quantified by absorbance at 260 nm, normalized to a uniform concentration, and supplemented with MgCl2 to a final concentration of 2.5 mM to support proper ncRNA folding prior to storage at −80 °C.

### Biochemical cDNA synthesis assays

ncRNAs were generated as described above and folded by heating to 95 °C and ramp cooling to 25 °C at a rate of 5 °C per minute. Reverse transcription reactions were then assembled by combining 2.5 µM RT and 2.5 µM ncRNA in Polymerization Buffer (50 mM Tris-HCl pH 8.0, 100 mM NaCl, 2 mM MgCl_2_, 5 mM TCEP) and incubating for 10 min at 37 °C to promote RT–ncRNA complex formation. cDNA synthesis was initiated by the addition of dNTPs, and reactions proceeded at 37 °C for 10 min unless otherwise indicated. Radioactive experiments typically contained 3.3 nM [α-^32^P]-dCTP (Revvity Health Sciences) together with an unlabeled (cold) dNTP mixture (dATP, dGTP, dTTP) at 100 µM each, added after complex formation. Reactions were terminated by heating to 95 °C for 2 min, prior to treatment with RNase A (Thermo Fisher Scientific) at 1 mg mL^-1^ and Proteinase K (Thermo Fisher Scientific) at 1 mg mL^-1^, as indicated. Samples treated with Proteinase K were incubated at 55 °C for 30 min. RNase A treatments were carried out at 37 °C for 30 min. Samples were boiled at 95 °C for 2 min between each post-reaction enzymatic treatment. For experiments comparing nucleotide incorporation across all four dNTPs (Fig. 3c), parallel reactions were assembled, each containing 3.3 nM of an individual [α-^32^P]-labeled dNTP. Reactions were terminated by heating at 95 °C for 2 min and then directly resolved by SDS-PAGE without further treatment◻

### Denaturing PAGE analysis of cDNA synthesis products

Radioactive DNA products from biochemical cDNA synthesis assays were resolved by 10% denaturing urea-PAGE. Samples were mixed in equal volumes with 2× RNA Loading Dye (95% formamide, 0.025% bromophenol blue, 0.025% SDS), denatured by heating at 95 °C for 2 min, and loaded onto gels for electrophoretic separation. Radioactive protein components were resolved by 12% denaturing SDS-PAGE. DNA ladders (20/100 nt from IDT and 1 kb+ from New England Biolabs) were 5′-radiolabeled using T4 PNK and [γ-^32^P]-ATP. Radiolabeling reactions (25 µL total) contained 100 µg/mL DNA ladder, 1× T4 PNK Reaction Buffer, 0.4 U µL◻¹ T4 PNK, and 33 nM [γ-^32^P]-ATP. Reactions were incubated at 37 °C for 30 min, heat-inactivated at 65 °C for 20 min, diluted to 50 µL with H2O, and purified using MicroSpin G-25 columns according to the manufacturer’s instructions, yielding 50 µg/mL 5′-radiolabeled DNA. Samples were mixed with 6× SDS Loading Dye (375 mM Tris-HCl pH 7.0, 9% SDS, 50% glycerol, 0.03% bromophenol blue), denatured by heating, and resolved by electrophoresis. For visualization, gels were dried for 2 h using a vacuum heat drier, exposed to a phosphor screen for 12 h, and imaged on a Typhoon imaging system (GE).

### Sequencing of biochemical cDNA synthesis reaction products

For sequencing-based analyses, biochemical cDNA synthesis reactions were assembled as described above using cold dNTPs at 100 µM each. Prior to downstream processing, a defined mixture of three 150-bp control oligonucleotides was spiked into each reaction at a uniform concentration to enable cross-sample normalization. DNA was isolated using the Monarch Spin PCR and DNA Cleanup kit (NEB), following the Oligonucleotide Cleanup protocol. Adapter ligation and conversion of ssDNA to dsDNA were performed using the xGen ssDNA & Low-Input DNA Library Prep Kit (IDT) with half the recommended reaction volumes. Libraries were sequenced on an Element AVITI in single-end mode with 150 cycles.

Sample reads were processed by Cutadapt (v5.0) to filter out adapter sequences, trim low-quality bases, and remove reads shorter than 15 bp, using a tiling approach across the 3′ adapter as previously described, with parameters -e 0.2, -O 10, -q 30, and -m 45. Trimmed reads were aligned with bwa-mem2 (v2.2.1) under default parameters to a reference containing the sequences of the three 150-bp spike-in oligonucleotides. Alignments were sorted and indexed using SAMtools (v1.17). Reads that failed to align were extracted using SAMtools fasta, and the fraction of unmapped reads was calculated using SAMtools flagstat. Motif-containing reads were quantified using a custom Python script: each read was screened for the expected tandem repeat motif, allowing a Hamming distance of 1 per repeat unit, and hits were normalized to spike-in-aligned reads to generate counts per million (CPM) values comparable across samples and sequencing runs. Motif position graphs were generated by identifying the first occurrence of the consensus motif in each read and tabulating subsequent motif occurrences at their corresponding downstream positions. 5× repeat reads were defined as those containing five tandem, uninterrupted copies of the motif.

### Cryo-EM Sample preparation and data collection

The RNA components of *Eco*_1_DRT10 and *Eco*_3_DRT10 RT-ncRNA were chemically synthesized (IDT) and folded according to previously established protocols^3^. The complexes were reconstituted by incubating purified RT proteins and folded ncRNA at a 1:1 molar ratio for 20 mins at 37 °C. The final reconstitution buffer was 20 mM HEPES pH 7.5, 300 mM NaCl, 7 mM MgCl_2_ for *Eco*_1_DRT10 and 20 mM HEPES pH 7.5, 350 mM NaCl, 8 mM MgCl_2_ for *Eco*_3_DRT10. Quanti-foil-R1.2/1.3 (300-mesh, gold) grids (Quantifoil Micro Tools) were used for *Eco*_1_DRT10 vitrification and UltrAufoil-R0.6/1.2 (300-mesh, gold) grids (Quantifoil Micro Tools) for *Eco*_3_DRT10. Grids were treated with a gas mix of 25/75% O_2_/Ar for 7s at 15W using a Gatan Solarus plasma machine (Gatan) prior to sample application A 3.5 µL aliquot of the samples were applied to the freshly plasma-treated grids, blotted for 4.5 s at 22°C and 95% relative humidity (blot force setting 0), and then vitrified by plunge-freezing into liquid ethane using a Vitrobot Mark IV (Thermo Fisher Scientific).

To capture the dNTP-bound state of the *Eco*_3_DRT10 RT-ncRNA complex, the complex was first reconstituted as described above in a buffer containing 20 mM HEPES pH 7.5, 500 mM NaCl, 9 mM MgCl_2_, and 0.0006% (w/v) lauryl maltose neopentyl glycol (LMNG). The dNTP-bound state was prepared via on-grid mixing: an aliquot of 1.8 µL of the pre-assembled RT-ncRNA complex was first applied to a freshly plasma-treated UltrAufoil R1.2/1.3 (300-mesh, gold) grid (Quantifoil Micro Tools), followed by the immediate addition of 1.8 µL of a 200 µM dNTPs mixture (prepared in 20 mM HEPES pH 7.5, 500 mM NaCl). Following brief on-grid mixing, the samples were immediately blotted for 4.5 s at 22 °C and 95% relative humidity (blot force 0) and then vitrified by plunge-freezing into liquid ethane using a Vitrobot Mark IV (Thermo Fisher Scientific).

Exploratory screening for particle density and ice quality was performed in a 200 KeV Glacios microscope equipped with a direct detector (Thermo Fisher Scientific) at the Columbia University Cryo-EM center and New York Structural Biology Center. Microscope operation and data collection were carried out using the Leginon software^42^. Best quality grids were selected for high resolution data collection.

For the *Eco*_1_DRT10 RT-ncRNA complex, a dataset of 25,980 movies was collected using the Leginon software on a Krios instrument (Thermo Fisher Scientific) at the New York Structural Biology Center. This instrument was equipped with a Cs corrector, a Gatan K3 direct electron detector integrated in a BioQuantum energy filter (using a 20-eV energy slit) (Gatan). The movies were acquired at a calibrated physical pixel size of 0.669 Å (after in-hardware binning), with a defocus range from -0.5 to -2.5 mm and a total dose of 53.63 e^−^/Å^−2^ (40 frames). The *Eco*_3_DRT10 RT-ncRNA complex dataset consisted of 11,765 movies and was also collected at the New York Structural Biology Center at a calibrated physical pixel size of 0.650 Å, with a defocus range from -0.2 to -1.5 mm and a total dose of 65.23 e^−^ Å^−2^ (60 frames). Finally, for the dNTP-bound *Eco*_3_DRT10 RT-ncRNA complex, 15,406 movies were collected on a Krios instrument (Thermo Fisher Scientific) using SerialEM^43^ at HHMI Janelia Research Campus. The dataset was acquired in super-resolution mode (pixel size 0.5305 Å) for a final binned calibrated physical pixel size of 1.061 Å, a defocus range from -0.8 to -2.0 mm and a total dose of 50 e^−^ Å^−2^ (50 frames). The detector in this instrument was a K3 (Gatan) also operated with a slit width of 20-eV.

### Cryo-EM data processing

*Eco*_1_DRT10 data was integrally processed in Relion3^44^ using a wrapper to ctffind4^45^ for the initial estimation of CTF values. The full dataset comprising 25,980 movies was motion corrected and the ctf values per micrograph were estimated using ctffind4. Particle picking was implemented first in a small subset of 100 motion corrected randomly selected images, initially using the Relion LOG picker^46^. Three iterative steps of c2d produced class averages of enough quality to train a topaz model that clearly overperformed the LOG picker in the small subset of images. The Topaz model^47^ was then used to pick the entire dataset producing more than 3 million hits. This large set of particles (extraction box 256 pixels, binned to 64 pixels) was submitted to 5 steps of c2d classifications using the classic Relion algorithm for c2d with increasing T parameter values (in increasing steps of 2) from a starting value of 4. This approach produced well defined multiple averages of a rigid-looking particle that in some views presented a pseudo-symmetric dimeric aspect. After c2d selection, the dataset was reduced to 654,888 particles which were submitted to a SGD step using 4 classes with a T value of 3. Only 1 of the 4 classes (comprising approximately half of the particles) produced a map that exhibited clear features compatible with known characteristics of proteins and nucleic acids. This subset of 335,549 particles was refined in 3D iteratively with increasing box sizes and performing local ctf-values estimations as well as local particle movement estimation using the bayeshian polishing approach in Relion^44^. The final map reaches a nominal resolution of approximately 3.2 Å showing excellent density as well as good angular distribution as there was no evidence of preferential orientation in the directionally homogenous map. The density for loop-A and B regions of the map was notably weaker compared to the rest of the map, suggesting flexible behaviour of these areas. Multiple 3D classification attempts using whole map approaches, masked approaches and local refinements were unsuccessful in localizing alternative conformation and/or in improving substantially the density in these areas. It is thus quite possible that both loops remain highly mobile as a requisite for function in the otherwise exceptionally rigid ribo-nucleic particle.

For model building, the refined unsharpened maps were post-processed in multiple ways to leverage the maximum amount of information. We used classic B-factor sharpened and masked post-processed maps from Relion, as well as deep-learning post-processed maps from DeepEMnhacer^48^, however, only the Relion sharpened map was used for stereochemical refinement. A single run of ModelAngelo^49^, using the known protein sequence of *Eco*_1_DRT10, was able to build roughly 90% of the protein component of the ribo-nucleic particle. We then used manual inspection in COOT^50^ to build and correct the rest of the protein residues. The RNA component was more challenging, ModelAngelo was unable to define a proper register for the RNA, we built it manually using a covariation model as guide. Notably, AlphaFold3^51^ predictions for the ribo-nucleic particle were wrong, unable to predict the dimeric architecture or the RNA fold. It however successfully predicted the overall fold of the protein monomer.

The final model comprising 121 nucleotides and 880 amino acids was subjected to a first round of stereochemical refinement using phenix.real_space_refinement with the secondary structure restraints option activated^52^. We then performed a final step of manual correction in COOT prior to a final step of stereochemical refinement in Fourier space using refemac5^53^ with secondary structure restraints calculated in ProSmart for proteins and LibG for the RNA component. Model validation was performed with Mol-probity

The dNTP-bound state and the apo-state of *Eco*_3_DRT10 RT-ncRNA datasets were processed using a similar image processing strategy. Movie frames were aligned for motion correction, and their contrast transfer function (CTF) parameters were estimated using the patch motion correction and patch CTF estimation algorithms in CryoSPARC^54^.

For the *Eco*_3_DRT10 RT-ncRNA complex dataset, a total of 10,382 micrographs remained after curation based on particle density and ice quality. Initially, particles were picked from 1,000 representative micrographs using the template-free blob picker and subjected to two rounds of 2D classification to remove non-particle artifacts and low-quality images. The resulting high-quality 2D classes served as templates for automated particle picking across the entire dataset. A total of 2,662,011 particles were picked and extracted from all micrographs at 2x binning (1.30 Å per pixel). The particles were then subjected to two rounds of 2D classification to remove unaligned particles, after which the selected particles (563,940 particles) were sorted into three reference-free 3D classes using *ab initio* modeling in CryoSPARC^54^. These three 3D classes were used as templates for two rounds of heterogeneous refinement, after which the selected particles (553,748) were re-extracted at the original pixel size (0.65 Å per pixel). These particles were subjected to two rounds of heterogeneous refinement to remove poor-quality particles. A subset of 271,876 particles was selected for 3D reconstruction using non-uniform refinement^55^. The particles were further refined with reference-based motion correction and one round of heterogeneous refinement. A final set of 233,422 particles yielded a reconstruction with a global resolution of 3.30 Å.

For the dNTP-bound state of the *Eco*_3_DRT10 RT-ncRNA complex dataset, a total of 14,487 micrographs were retained after curation based on particle density and ice quality. To generate high-quality templates for automated picking and subsequent heterogeneous refinement, a representative subset of 5,000 micrographs was initially processed. Particles picked via the template-free blob-picker were subjected to three rounds of 2D classification to remove non-particle artifacts. The resulting clean 2D class averages were then selected as templates for comprehensive automated particle picking across all 14,487 micrographs. To establish reliable 3D references, 471,382 high-quality particles selected from the initial 5,000-micrograph subset were sorted into four reference-free 3D classes using *ab initio* modeling in CryoSPARC. Three of these four classes were subsequently employed as initial volumes for the heterogeneous refinement of the full dataset. For the entire dataset, a total of 10,443,595 particles were picked and extracted from all micrographs at 2x binning (2.122 Å per pixel). These particles are subjected to two rounds of heterogeneous refinement using the aforementioned *ab initio* models as templates. A subset of 3,736,051 selected particles was then re-extracted at the original pixel size (1.061 Å/pixel) and further pruned through two additional rounds of heterogeneous refinement to eliminate poor-quality particles and resolve structural heterogeneity. A total of 1,957,058 particles were selected for high-resolution 3D reconstruction using the non-uniform refinement in CryoSPARC. After reference-based motion correction and one additional round of heterogeneous refinement, a final set of 1,022,101 particles was selected for the final non-uniform refinement, yielding a reconstruction with a global resolution of 2.97 Å.

### DRT10 Bioinformatics

To assess conservation of DRT10 RT C-termini, we analyzed a previously described multiple sequence alignment of DRT10 RT sequences^3^. Unaligned sequences were scanned with PSIPRED^56^ in single-sequence mode to classify each residue into predicted coil, alpha-helix, or beta-strand states. Residues were then renumbered with their corresponding positions in the multiple sequence alignment. To assess core conserved protein content, residues with less than 50% conservation in the multiple sequence alignment were removed. Secondary structure predictions for the remaining RT residues were then plotted, alongside the total number of proteins with occupancy for each remaining position in the trimmed multiple sequence alignment. The C-terminal α-helix, mapped relative to the *Eco*3 reference sequence (OKX54664.1), was explicitly annotated to assess its structural conservation. Data integration, filtering, and graphical visualizations were executed in R utilizing the *Biostrings* (v2.8.0) and *tidyverse* (v2.0.0) packages.

To examine the pseudo-symmetric nature of DRT10 ncRNAs, secondary structures were extracted from the *Eco*_1_*/Eco*_3_ structures or AlphaFold 3^51^ predictions of two copies of the DRT10 RT and one copy of the cognate ncRNA. Dot-bracket annotations were extracted using the DSSR web application^57^. Predicted structures were manually refined via comparisons to predictions made with RNAfold from the Vienna RNA package^58^, prior to visualization with VARNA^59^.

## Data availability

Next-generation sequencing data will be made available in the National Center for Biotechnology Information (NCBI) Sequence Read Archive (BioProject accession: PRJNA1470254) at the time of publication. The Cryo-EM maps were deposited to the Electron Microscopy Data Bank under the accession codes EMD-57959 (*Eco*_1_DRT10 apo complex), EMD-57969 (*Eco*_3_DRT10 apo complex) and EMD-57972 (*Eco*_3_DRT10 dNTP bound complex). The corresponding atomic coordinates were deposited in the Protein Data Bank under the accession code 30QM (*Eco*_1_DRT10 apo complex), 30QU (*Eco*_3_DRT10 apo complex) and 30QX (*Eco*_3_DRT10 dNTP bound complex). Datasets generated and analyzed in the current study are available from the corresponding authors on reasonable request.

## Code availability

Custom scripts used for bioinformatics, *in vitro* transcription sequencing data analyses are available upon request.

## ACKNOWLEDGMENTS

We thank T.M. Smith, A.J. Robinson, and R. Rafat for laboratory support; O.B. Clarke and A.W.P. Fitzpatrick for their generous help with cryo-EM grids; the staff scientists at the Simons Electron Microscopy Center at the New York Structural Biology Center with microscope operation and data collection; the staff scientists at HHMI Janelia CryoEM Facility for help with microscope operation and data collection; the JP Sulzberger Columbia Genome Center for NGS support; and M. Wang, R. Žedaveinytė, and all members of the Sternberg Lab for valuable discussions.

## AUTHOR CONTRIBUTIONS

J.L.R., Q.H., I.S.F., and S.H.S. conceived the project. J.L.R. isolated the RT–ncRNA complex, together with Q.H., and performed all biochemical and high-throughput sequencing assays. Q.H. prepared cryo-EM grids and collected cryo-EM data, together with J.W., and tested protein priming mutants. T.W. performed bioinformatics analyses and assisted in figure design. G.D.L. contributed to experimental design and data interpretation. S.T. performed phylogenetic analyses and assisted with experimental design and data visualization. I.S.F. analyzed cryo-EM data and built atomic models, together with Q.H. I.S.F. and S.H.S. supervised the study. J.L.R., Q.H., I.S.F., and S.H.S. discussed the data and wrote the manuscript, with input from all authors.

## COMPETING INTERESTS

Columbia University has filed a patent application related to this work. S.H.S. is a co-founder and scientific advisor to Dahlia Biosciences, a scientific advisor to CrisprBits and Prime Medicine, and an equity holder in Dahlia Biosciences and CrisprBits. The remaining authors declare no competing interests.

Correspondence and requests for materials should be addressed to I.S.F. or S.H.S.

