## Supplementary Figures for "Mechanism of tandem-repeat DNA synthesis by an antiviral reverse transcriptase"

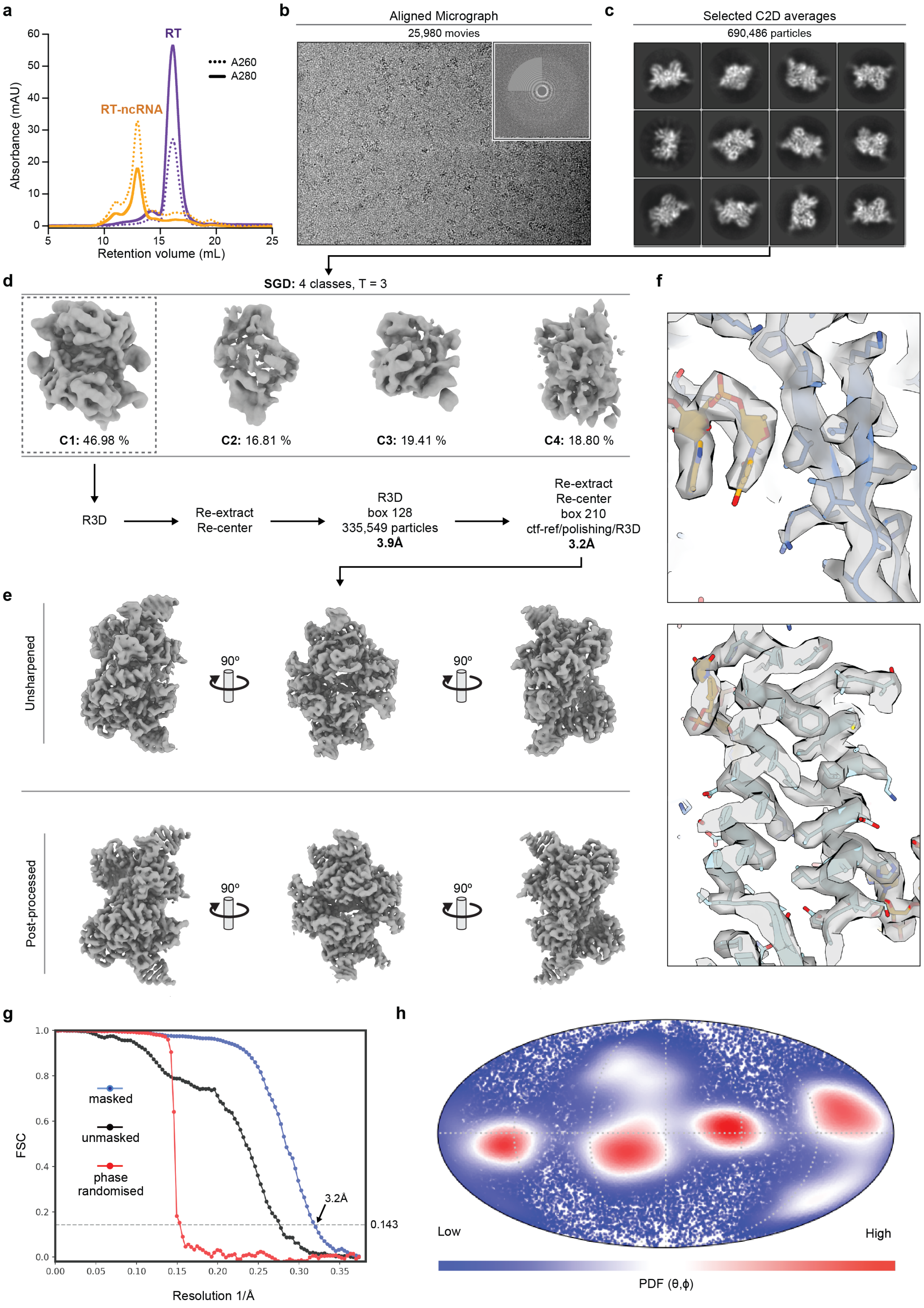

**Figure S1 | Cryo-EM data processing and validation for the *Eco*$_1$DRT10 RT–ncRNA complex. a,** Size-exclusion chromatography profiles of *Eco*$_1$DRT10 RT in the absence and presence of ncRNA, on a Superdex 200 Increase 10/300 column, showing a shift in elution volume and increased A260/A280 upon complex formation. **b,** Representative motion-corrected cryo-EM micrograph from 25,980 movies for the *Eco*$_1$DRT10 RT–ncRNA complex. **c,** Selected 2D class averages. **d,** Cryo-EM data processing workflow. Stochastic Gradient Descent (SGD) analysis of the particles selected after recursive C2D runs yielded four classes, with class 1 (~47% of particles) exhibiting features compatible with known protein and RNA motifs, including helices and clear domains, used for 3D refinement. Iterative re-extraction, re-centering, CTF refinement, and Bayesian polishing improved the resolution to 3.2 Å. **e,** Final reconstruction shown in multiple orientation, in both unsharpened (top) and post-processed (bottom) form. **f,** Representative cryo-EM density overlaid with the final, stereochemically refined model. **g,** Fourier Shell Correlation (FSC) curves for the final reconstruction, indicating a global resolution of 3.2 Å. **h,** Mollweide spherical projection of the Eulerian angles for all particles obtained after the final refinement.

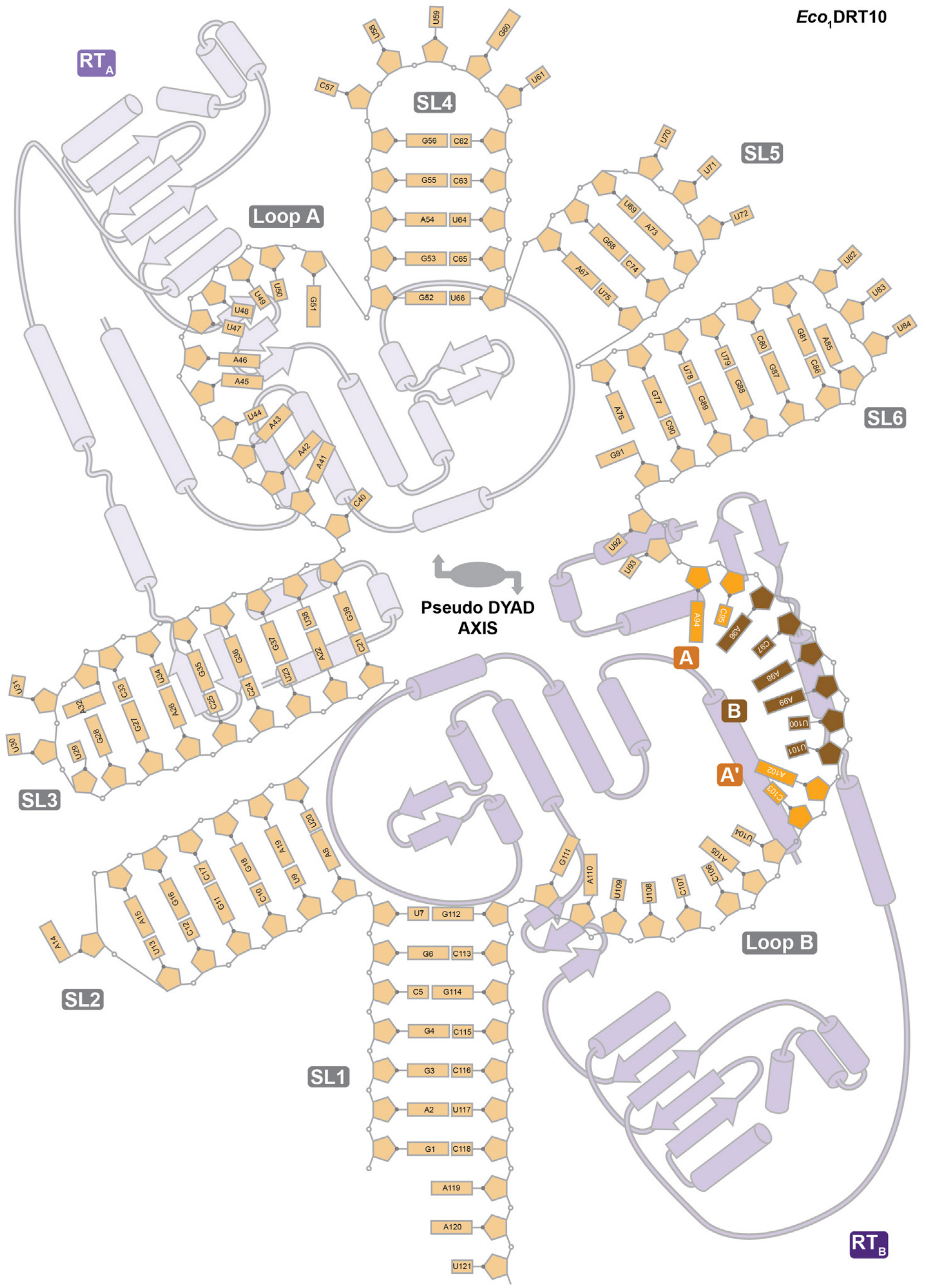

**Figure S2 | Secondary structure and pseudo-symmetric organization of the $Eco_1$DRT10 ncRNA.** Annotated secondary structure of the ncRNA highlights stem-loop elements (SL1–SL6), Loop A and Loop B, and the internal template-containing region. Nucleotide identities are shown, and base-pairing interactions are indicated. The ncRNA adopts an overall pseudo-symmetric architecture organized around a central dyad axis, with two structurally related halves that engage the RT dimer (monomers A and B). Regions contacting each RT promoter are indicated, illustrating how the RNA scaffold bridges the two subunits and organizes the complex.

a

b

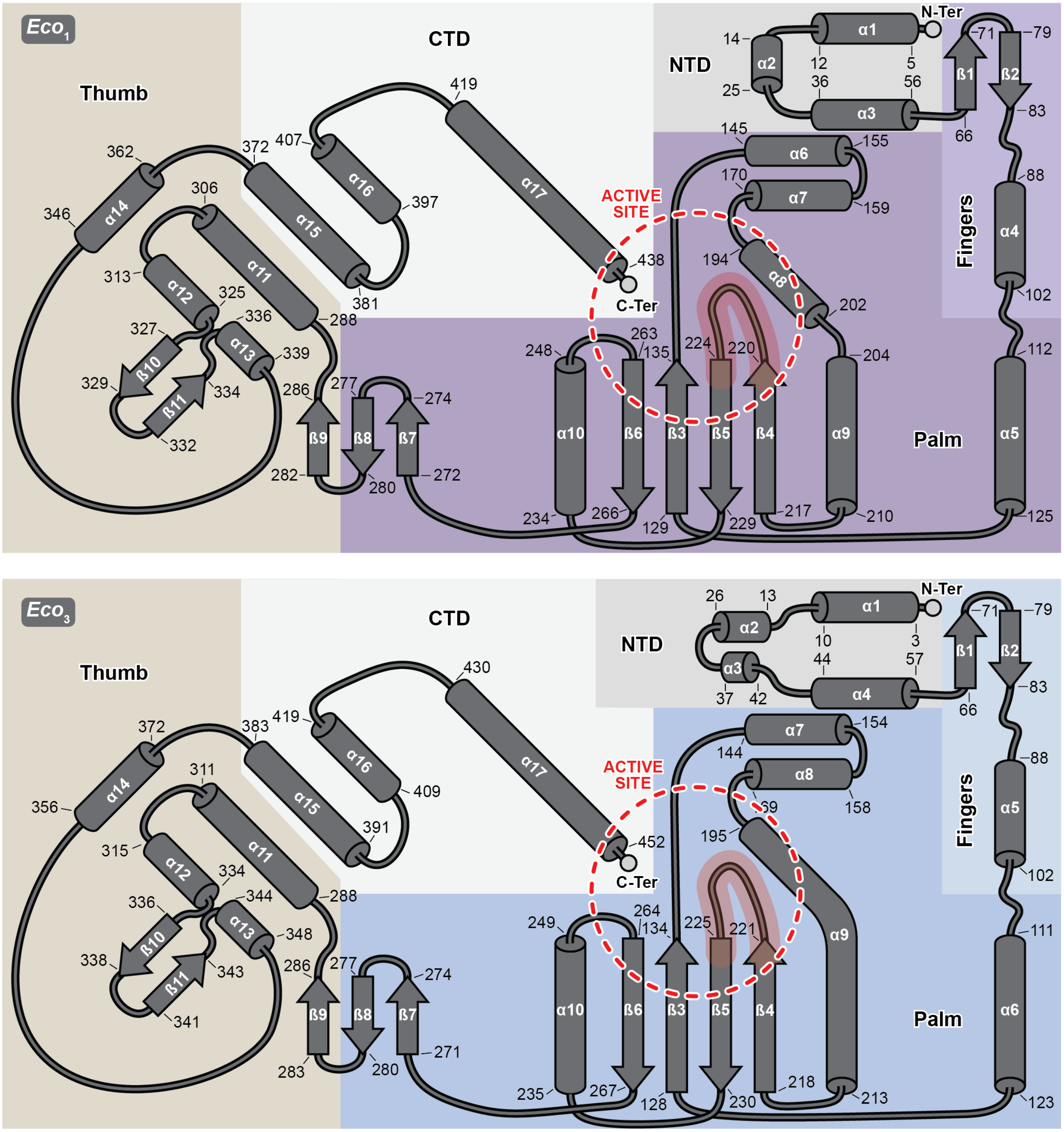

**Figure S3 | Secondary structure of the *Eco*$_1$DRT10 and *Eco*$_3$DRT10 RT.** Annotated secondary structure of the RT from *Eco*$_1$DRT10 (**a**) and *Eco*$_3$DRT10 (**b**).

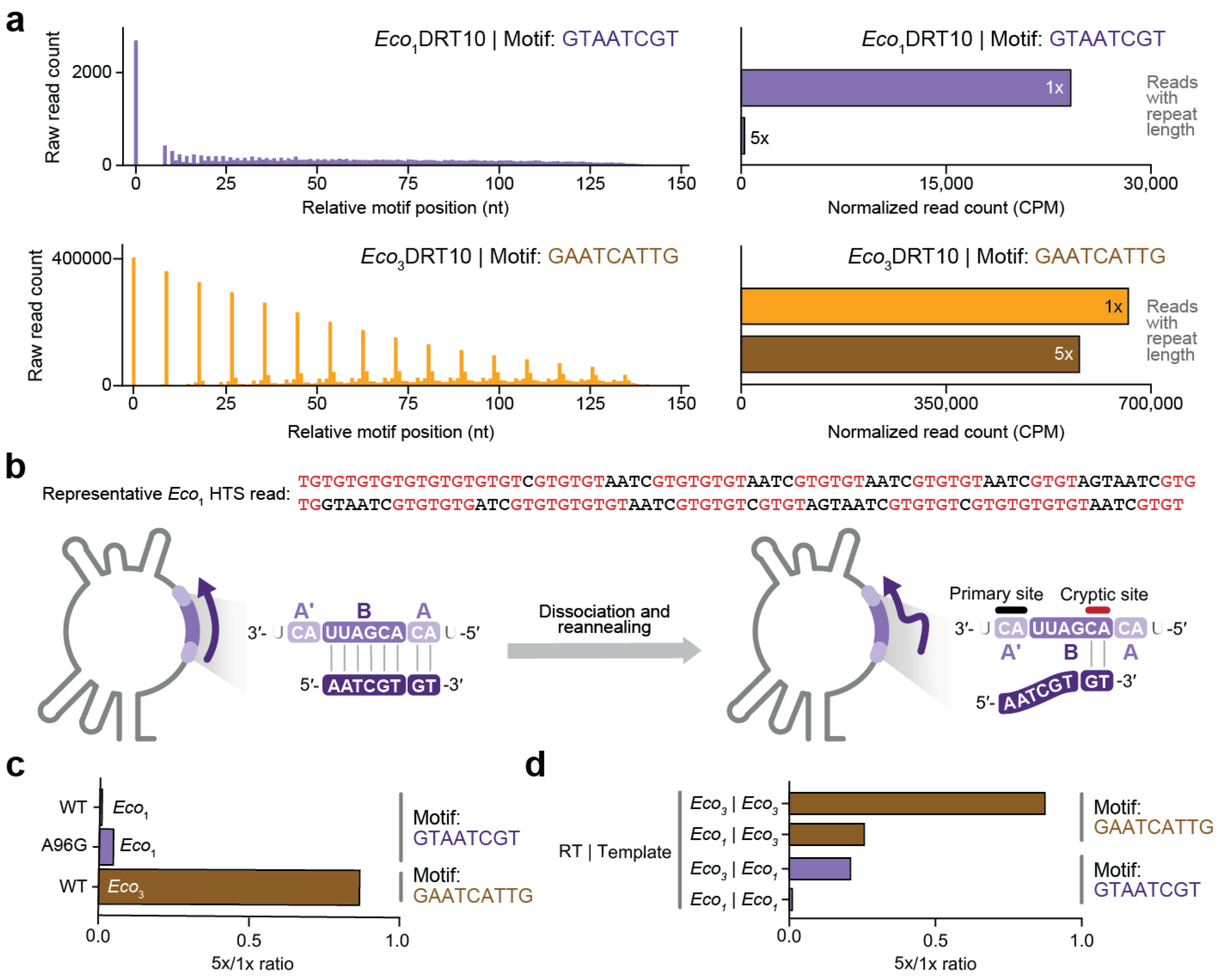

**Figure S4 | Characterization of *Eco*$_1$DRT10 and *Eco*$_3$DRT10 cDNA synthesis products and repeat addition fidelity. a,** High-throughput sequencing (HTS) analysis of *Eco*$_1$ and *Eco*$_3$ cDNA synthesis products, visualized using two complementary methods. *Left*, motif heat maps, in which the first occurrence of the consensus motif in a given read defines the start position, and subsequent motif occurrences throughout the read are counted at their corresponding positions. *Right*, bar graph quantification of reads containing 1× or 5× perfect repeats, normalized by counts per million (CPM) using a control oligonucleotide spike-in added uniformly across sequencing runs. **b,** *Top*, representative sequencing analysis showing aberrant GT-repeat tracts (red) resulting from cryptic-site resetting during cDNA synthesis. *Bottom*, schematic of the *Eco*$_1$ ncRNA template region highlighting the cryptic internal reannealing motif responsible for this resetting. **c,** Sequencing analysis of an *Eco*$_1$ ncRNA variant (A96G) designed to disrupt the cryptic repeat, showing mild restoration of accurate repeat synthesis. Data are presented as the ratio of reads containing a 1× repeat to those containing a 5× repeat, in order to isolate RAP accuracy from overall cDNA abundance. **d,** Template-swapping experiments between *Eco*$_1$ and *Eco*$_3$ systems, assessing the contribution of RT versus template sequence to repeat synthesis fidelity. Introduction of the *Eco*$_1$ template into the *Eco*$_3$ ncRNA increases RAP accuracy, whereas the *Eco*$_3$ template in the *Eco*$_1$ ncRNA reduces it. Data are presented as the ratio of reads containing a 1× repeat to those containing a 5× repeat. Data shown in **a, c,** was repeated twice with similar results.

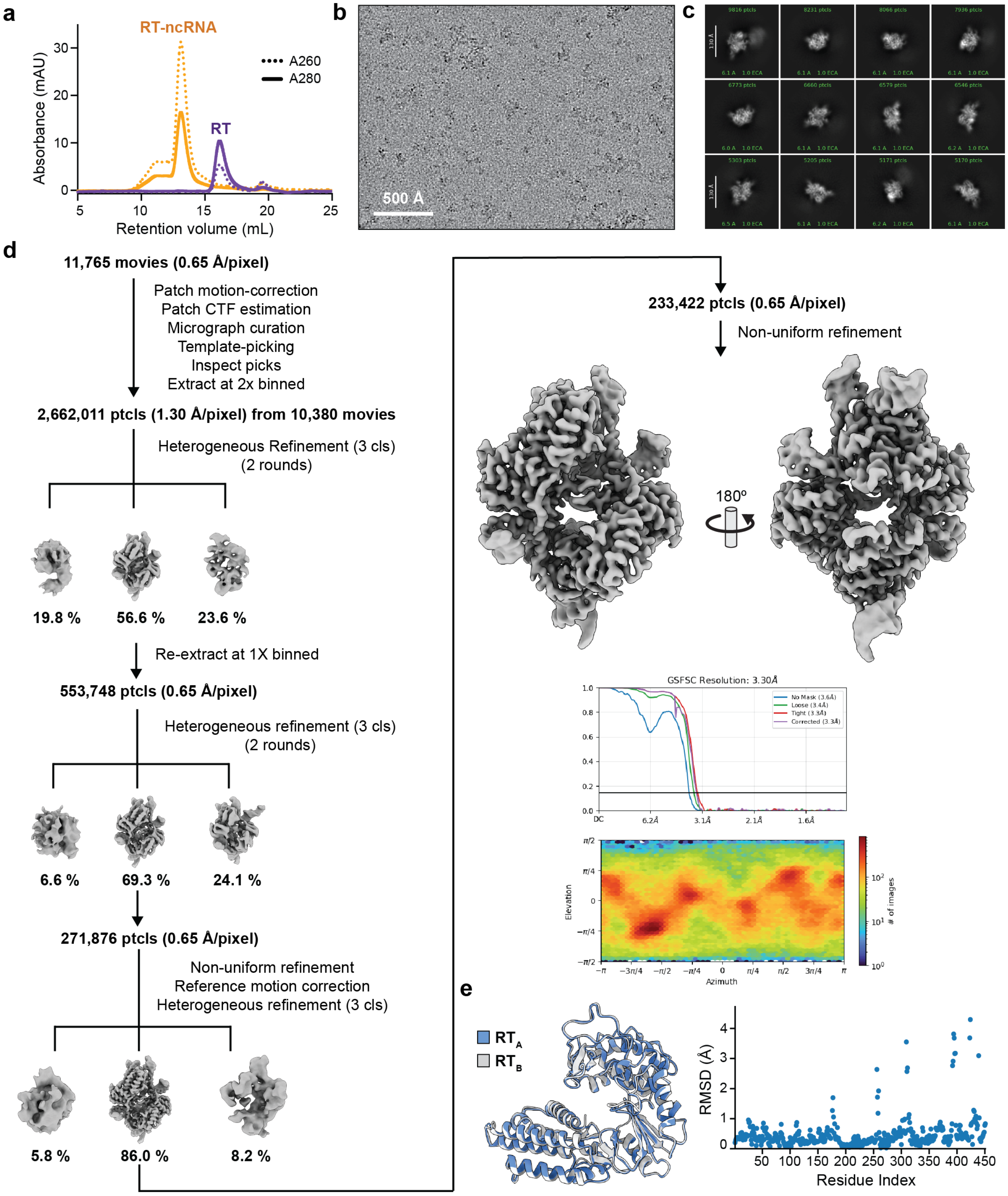

**Figure S5 | Image processing workflow for the *Eco*$_3$DRT10 RT-ncRNA complex. a,** Size-exclusion chromatography profiles of *Eco*$_3$DRT10 RT in the absence and presence of ncRNA, on a Superdex 200 Increase 10/300 column, showing a shift in elution volume and increased A260/A280 upon complex formation. **b,** Representative motion-corrected cryo-EM micrograph from 11,765 movies for the *Eco*$_1$DRT10 RT–ncRNA complex. The scale bar dimension is 500 Å. **c,** Selected 2D class averages. A 130 Å scale bar is shown in white. **d,** Cryo-EM data processing workflow employed to obtain the consensus map of the *Eco*$_3$DRT10 RT-ncRNA complex. **e,** Both *Eco*$_3$RT monomers showing nearly indistinguishable conformations with equivalent domain organization, with a RMSD of 0.725 Å over 448 aligned Cα atoms.

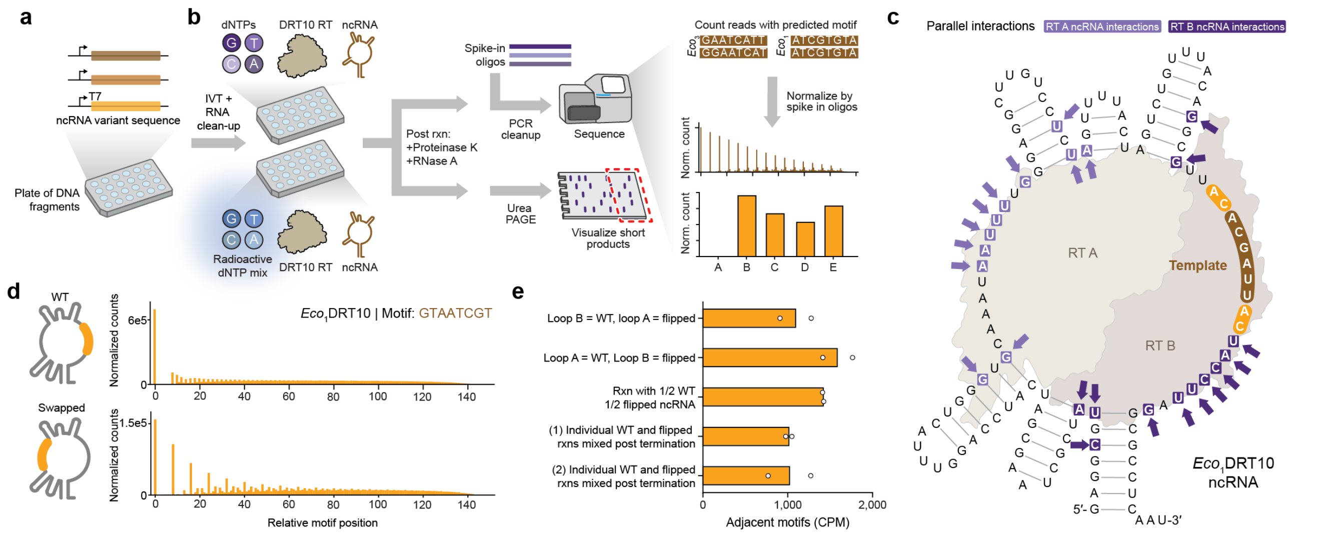

**Figure S6 | *In vitro* pipeline for ncRNA variant generation and cDNA product evaluation and $Eco_1$ symmetry. a,** Schematic of ncRNA variant production: DNA fragments arrayed in a 96-well plate serve as templates for *in vitro* transcription, followed by RNA cleanup (Zymo kit). **b,** Schematic of *in vitro* cDNA synthesis and product analysis: ncRNA variants are incubated with the DRT10 RT and dNTPs, followed by Proteinase K and RNase A treatment. For sequencing-based analysis of long products, a control oligonucleotide of known sequence and quantity is spiked in for downstream CPM normalization prior to PCR cleanup, library preparation, and NGS. Processed reads are analyzed for repeats of the expected template motif. Alternatively, products are resolved by radiolabeled denaturing urea-PAGE, which enables visualization of short products that are lost in sequencing-based workflows. **c,** Cartoon schematic of the $Eco_1$DRT10 RT–ncRNA complex highlighting the putative line of symmetry and symmetric RT–ncRNA interactions on opposite sides of the ncRNA. **d.** Diagram of the $Eco_1$ ncRNA loop-swapping ncRNA (left), with accompanying *in vitro* cDNA sequencing results shown as motif heat maps, demonstrating that the two loops can be functionally exchanged (right). Data shown was repeated twice with similar results. **e,** Sequencing-based assay distinguishing intermolecular from intramolecular template usage. Reactions contained either two distinct ncRNAs co-incubated in a single tube (intermolecular template usage) or a single ncRNA carrying a unique template in each of loops A and B (intramolecular template usage). Reads spanning the junction formed by the two distinct motifs were counted in each case and compared to a negative control in which two separate single-ncRNA reactions were terminated and then mixed (two biological replicates). Junction counts are reported as reads per million, normalized to a spike-in oligonucleotide. Data are mean of $n=2$ independent replicates.

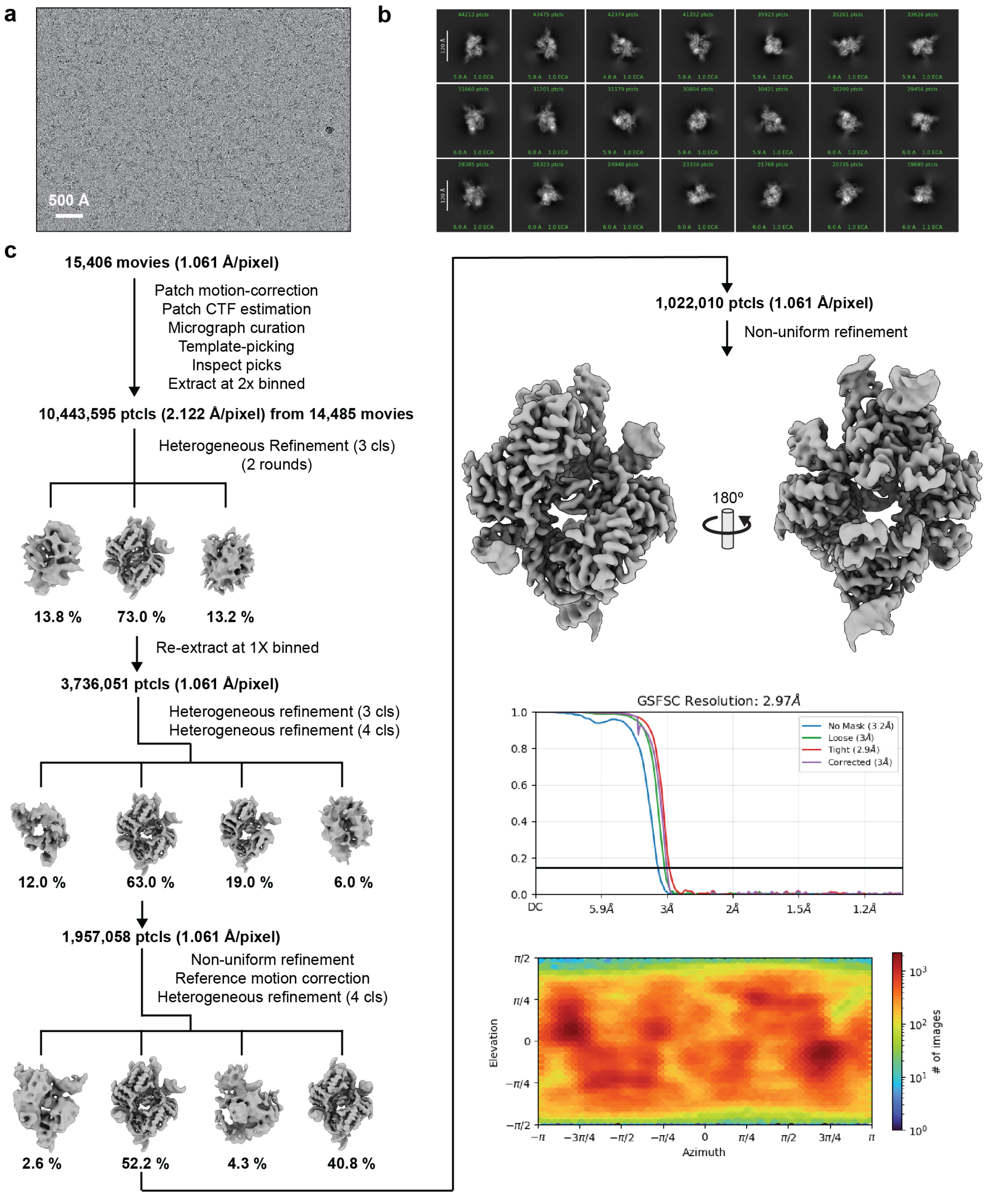

**Figure S7 | Image processing workflow for the dNTP-bound *Eco*$_3$DRT10 RT-ncRNA complex. a,** Representative motion-corrected cryo-EM micrograph from 15,406 movies for the dNTP-bound *Eco*$_3$DRT10 RT–ncRNA complex. The scale bar dimension is 500 Å. **b,** Selected 2D class averages. A 120 Å scale bar is shown in white. **c,** Cryo-EM data processing workflow employed to obtain the consensus map of the *Eco*$_3$DRT10 RT-ncRNA complex.

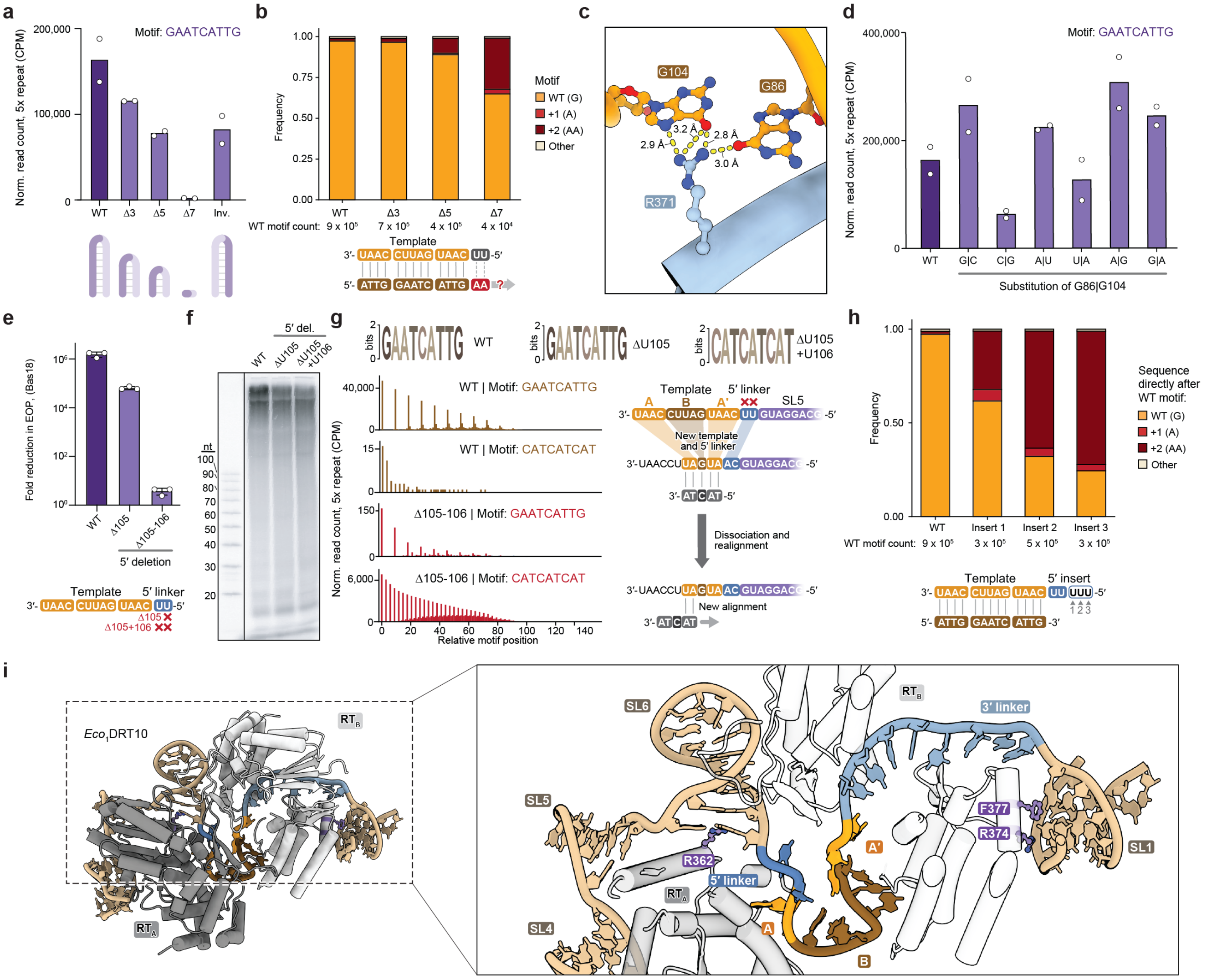

**Figure S8 | Structural and biochemical characterization of SL5 template boundary determinants. a,** High-throughput sequencing analysis of cDNA products from progressive SL5 base-pair truncations, normalized by CPM using a spike-in control oligonucleotide. **b,** Analysis of template read-through in SL5 truncation variants. After identifying reads containing the WT motif (GAATCATTG), the sequence immediately following the motif was analyzed, quantified as the frequency of each nucleotide identity at positions downstream of the WT motif boundary. **c,** Structural zoom-in on the non-canonical GG Hoogsteen conformation (G104–G86) at the base of SL5, showing the bidentate hydrogen-bonding interaction with R371. **d,** High-throughput sequencing analysis of cDNA products from G104–G86 base-pair identity swaps, normalized by CPM using a spike-in control oligonucleotide. **e,** *In vivo* phage defense assays for progressive nucleotide deletions in the 5' linker region, showing loss of defense with increasing truncation. Data are mean ± s.d. of $n=3$ technical replicates. **f,** Radiolabeled ([α-$^{32}$P]-dCTP) denaturing PAGE analysis of 5' linker truncation variants, revealing apparent production of long-length product even in cases previously identified by sequencing as failing to produce WT motif repeats. **g,** Sequencing analysis of 5' linker truncation variants. *Top*, MEME motif analysis showing retention of the WT motif in the Δ1 nt variant and emergence of a novel CAT repeat motif in the Δ2 nt variant. *Bottom*, sequencing motif graphs looking for WT and novel motifs. *Bottom right*, cartoon schematic illustrating a putative alternative RAP strategy in which the RT stops 2 nt before the WT template boundary, utilizing a new AT reannealing site to sustain processive synthesis of the aberrant repeat product. **h,** Analysis of template read-through in 5' linker insertions. Reads containing the WT motif are identified and the sequence immediately following the motif is analyzed, quantified as the frequency of each nucleotide identity at positions 2 nt downstream of the WT motif boundary. **i,** Structural view of the template-boundary region in the $Eco_1$ complex, with the 5′ and 3′ linkers and their anchoring stem-loop contacts highlighted. For **a, d,** data are mean of $n=2$ independent replicates. Data shown in **b, f, g, h,** was repeated twice with similar results.

### SUPPLEMENTARY TABLES

Supplementary Table 1 | Description and sequence of plasmids used in this study.

Supplementary Table 2 | Strains used in this study.

Supplementary Table 3 | DRT10 proteins tested in this study.

Supplementary Table 4 | Oligonucleotides used in this study.

Supplementary Table 5 | Cryo-EM data collection, refinement, and validation statistics.

Supplementary Table 6 | DRT10 ncRNA variants tested in this study.

Supplementary Table 7 | DRT10 alternate template motifs.

**Supplementary Table 5 | Cryo-EM data collection, refinement, and validation statistics.**

| | ***Eco*** $_{1}$**DRT10 apo complex map** (EMD-57959) (PDB 30QM) | ***Eco*** $_{3}$**DRT10 apo complex map** (EMD-57969) (PDB 30QU) | ***Eco*** $_{3}$**DRT10 dNTP bound complex map** (EMD-57972) (PDB 30QX) |
|---|---|---|---|
| **Data collection and processing** | | | |
| Magnification | 105,000x | 130,000x | 81,000x |
| Voltage (kV) | 300 | 300 | 300 |
| Electron exposure (e–/Å$^2$) | 53.63 | 65.23 | 50 |
| Defocus range (μm) | -0.5 to -2.5 | -0.2 to -1.5 | -0.8 to -2.0 |
| Pixel size (Å) | 0.669 | 0.65 | 1.061 |
| Symmetry imposed | C1 | C1 | C1 |
| Initial particle images (no.) | 1,234,644 | 2,662,090 | 15,976,791 |
| Final particle images (no.) | 335,549 | 233,422 | 1,022,010 |
| Map resolution (Å) | 3.2 | 3.3 | 3 |
| FSC threshold | 0.143 | 0.143 | 0.143 |
| **Refinement** | | | |
| Initial model used (PDB code) | ModelAngelo/ Manual build | ModelAngelo/ Manual Build | ModelAngelo/ Manual Build |
| Model resolution (Å) | 3.5 | 3.5 | 3.5 |
| FSC threshold | 0.5 | 0.5 | 0.5 |
| Model resolution range (Å) | 3.3-5 | 3.3-5 | 3.3-5 |
| Map sharpening *B* factor (Å$^2$) | -89 | -77 | -69 |
| Q-Score | 0.477 | 0.494 | 0.497 |
| Model composition | | | |
| Non-hydrogen atoms | 9821 | 9913 | 9975 |
| Protein residues | 879 | 900 | 900 |
| Ligands | 0 | 0 | 2 |
| *B* factors (Å$^2$) | | | |
| Protein | 94.35 | 33.1 | 38.8 |
| Nucleotide | 115.9 | 86.36 | 117.2 |
| Ligand | N/A | N/A | 65.5 |
| R.m.s. deviations | | | |
| Bond lengths (Å) | 0.0091 | 0.0021 | 0.0037 |
| Bond angles (°) | 1.63 | 0.49 | 0.77 |
| Validation | | | |
| MolProbity score | 2.1 | 1.38 | 1.5 |
| Clashscore | 6.01 | 4.04 | 4.02 |
| Poor rotamers (%) | 2.01 | 0.37 | 1.1 |
| Ramachandran plot | | | |
| Favored (%) | 93.5 | 96.5 | 93.8 |
| Allowed (%) | 5.8 | 3.3 | 5.7 |
| Disallowed (%) | 0.7 | 0.2 | 0.5 |
